# Prevalence of Colorectal Cancer Molecular Profiles Is Not Captured by a Single Age Threshold

**DOI:** 10.64898/2026.08.13.743895

**Authors:** Luis Bermúdez-Guzmán, Iannish Sadien, MB BChir, Allan Ramos-Esquivel, Warner Alpízar-Alpízar

## Abstract

Early- and average-onset colorectal cancer (CRC) are separated at age 50, but whether this defines a biological threshold remains unclear. To clarify this, we identified molecular profiles in nine harmonised cBioPortal CRC cohorts (4,609 patients) by fitting Bernoulli mixture models to 31 repair-state, genomic-burden and gene-alteration features, excluding age, sex and tumour site, and compared their prevalence using <50/≥50 and decade-resolved groups. Four profiles captured conventional/CIN-like (P1), intermediate MSS (P2), KRAS/PI3K/APC-rich (P3) and hypermutated/MSI-high (P4) states along a left-to-right anatomical gradient. Although molecular identities remained stable, profile prevalence followed non-linear P1/P4 and opposing linear P2/P3 age trajectories. Profile prevalence patterns did not follow age distance: profile composition at 30-39 differed from 50-59 but not clearly from 40-49 or 60-69. The age-50 threshold captured only 17.8% of decade-resolved deviance, whereas the optimal age-70 cut-off retained only 51.3%. Validation in 2,235 non-overlapping MSK-IMPACT patients (2,134 age-evaluable) reproduced molecular-feature patterns (r=0.96-1.00), age trajectories (r=0.90) and limited binary-threshold performance: age 50 and the optimal age-75 cut-off retained 6.2% and 49.4%, respectively. Thus, age reorganises the prevalence of shared CRC states rather than defining a biological threshold at age 50.

## Main

CRC is biologically diverse, and this diversity unfolds across the colorectum and life course through differences in risk, precursor pathways, molecular profiles and clinical outcomes^1^. This diversity reflects recurrent genomic routes, including chromosomal instability, hypermutation most often associated with mismatch-repair deficiency and microsatellite instability, and serrated neoplasia^2,3^. Broader transcriptional states are captured by the consensus molecular subtype (CMS) framework, which integrates tumour-cell programmes with immune, stromal, metabolic and mesenchymal features^4^. These states are not distributed uniformly: anatomy and sex further shape them, with right- and left-sided CRC differing in mismatch repair status, BRAF mutation frequency, chromosomal instability, microbiome context, clinical outcome and therapeutic approach ^2,5–7^.

On top of these anatomical and sex-specific differences, variation across age at diagnosis adds a dimension that is commonly reduced to the early-onset versus average-onset distinction at 50 years. Although this binary threshold is particularly useful for tracking rising CRC incidence in younger generations^8–12^, it remains unclear whether it captures a discrete change in molecular-profile prevalence. In that sense, dichotomising age can obscure differences among decades within each group because it tests only the average difference between patients below and above the cut-off. In practice, associations with age at diagnosis may reflect life-course ageing, birth cohort, exposome, calendar period, screening and cohort ascertainment. Recent work on rising early-onset cancers has therefore called for molecularly defined frameworks that can connect epidemiological change to underlying tumour biology^13^. The unresolved question is therefore whether age-defined groups contain unique molecular states or differ in the prevalence and context of shared ones.

To address this question, we rebuilt a harmonised patient-level dataset of 4,609 primary CRCs and fitted Bernoulli mixture models (BMMs) to 31 broadly available genomic, repair-status and burden features. Age, sex and tumour site were withheld from model training and introduced only after profile discovery. Crucially, we uncoupled two frequently conflated parameters: preservation of molecular profile identity across age and redistribution of profile prevalence across age. Decade-resolved analysis uncovered non-linear P1/P4 prevalence trajectories and opposing linear P2/P3 trends, whereas the age-50 split captured only 17.8% of the profile-composition deviance resolved across decades. Validating these dynamics, the same molecular identities unfolded in a non-overlapping MSK-IMPACT validation cohort, in which the age 50 binary split retained only 6.2% of decade-resolved deviance. Ultimately, these findings redefine age-associated CRC biology as the redistribution of shared molecular states rather than a threshold between two different molecular diseases.

## Results

### A four-profile Bernoulli mixture model resolves the main CRC molecular states

The discovery dataset comprised 4,609 primary CRC patients from nine harmonised cBioPortal cohorts (Supplementary Table 1). Age at diagnosis was available for 4,607 patients (median 59 years, IQR 47-69), including 1,462 patients diagnosed before age 50 (31.7%). Among patients with known sex (n=4,556), 2,443 were male and 2,113 were female. Tumour site was available for 4,356 patients, including 1,865 left/distal colon, 1,247 right/proximal colon and 1,244 rectal cancers. Stage at diagnosis was available for 4,100 patients; race and ethnicity annotations were available for 3,298 and 720 patients, respectively (Supplementary Table 2).

To identify molecular profiles independently of age, sex, and anatomy, we fitted Bernoulli mixture models (BMMs) to 31 binary molecular features while withholding those three variables. The feature set was restricted to established CRC repair-state, genomic-burden and driver/pathway markers that could be harmonised as binary variables across both whole-exome and targeted-panel cohorts. K=4 (number of different molecular profiles, P1-P4) was selected as the principal resolution because it best balanced model fit, posterior classification confidence, profile size and biological interpretability (Figure 1A; Supplementary Table 3). Each of the nine retained cohorts contained a mixture of profiles, with no profile reducible to a single study (Supplementary Figure 1). P1 consisted of a conventional/CIN-like profile enriched for *APC* and *TP53* alterations, microsatellite stable/mismatch repair-proficient (MSS/pMMR), and high fraction of the genome altered (FGA-high) status (n=1,859). P2 was an intermediate/mixed MSS profile with frequent *TP53* alteration and FGA-high status but very low *APC* mutation burden (n=1,161). P3 showed an MSS KRAS/PI3K/APC-rich profile (n=1,009). P4 resolved as a profile enriched for microsatellite instability-high/deficient mismatch repair (MSI-H/dMMR), high tumour mutational burden (TMB-high; ≥10 mutations/Mb) status, *BRAF and RNF43 alterations* (n=580). The complete 31-feature prevalence matrix for P1-P4 is provided in Supplementary Table 4.

**Figure 1.**
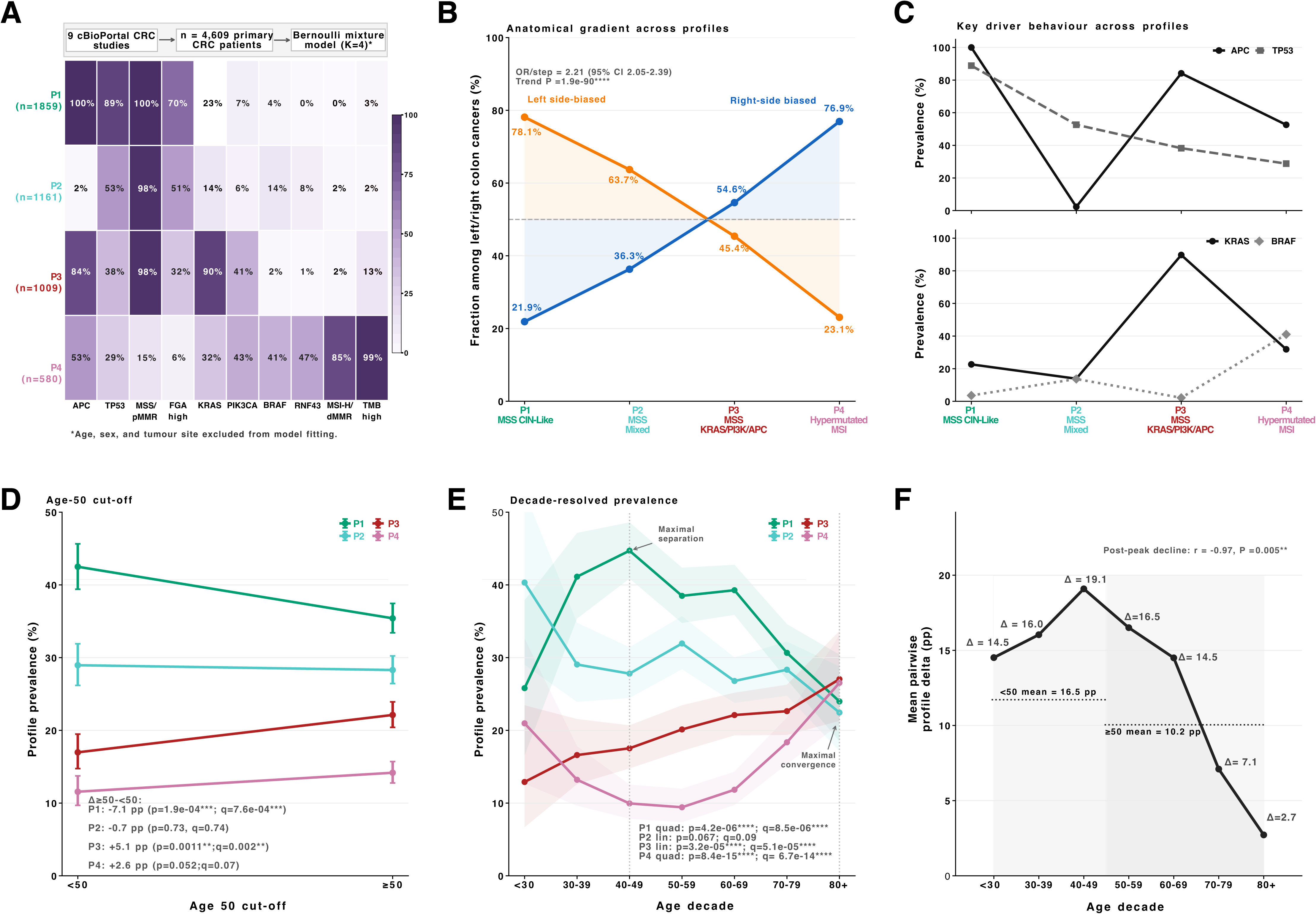
Profile discovery and decade-resolved age non-linearity. A. BMM K=4 prevalence heatmap of selected profile-defining molecular features and study/patient/model schematic; the complete 31-feature matrix is provided in Supplementary Table 4. Age, sex and tumour site were excluded from model fitting. B. Anatomical gradient of P1-P4 among left/right colon cancers. C. Key driver-gene alteration behaviour across profiles. D. Conventional <50 versus ≥50 profile prevalence comparison. E. Decade-resolved profile prevalence with selected trend statistics. F. Mean pairwise profile delta across age decades.

Despite tumour site being excluded from model training, the profiles mapped onto a strong left-to-right anatomical gradient (Figure 1B). P1 was predominantly left/distal-enriched and P4 strongly right/proximal-enriched; P2 and P3 occupied intermediate positions, with P2 shifted to the left and P3 shifted towards the right colon. The odds ratio for right-sided location was 2.21 per ordered profile step (95% CI 2.05-2.39; trend p=1.9×10^-90^). Figure 1C highlights the APC/TP53, KRAS and BRAF contrasts underlying these axes. Although our profiles were not built on transcriptomic data, they also aligned with established genomic components of the CMS framework. Across eight shared molecular features, P1 was closest to CMS2 (Pearson r=0.935; q=0.005), P3 to CMS3 (r=0.892; q=0.012) and P4 to CMS1 (r=0.937; q=0.005), whereas P2 showed partial proximity to CMS4 (r=0.662; q=0.168; Supplementary Figure 2).

### The age-50 threshold compresses decade-resolved variation in molecular-profile prevalence

We next examined whether the epidemiological age-50 threshold aligns with distinct tumour biology by evaluating profile composition across early-versus average-onset CRC. P1 prevalence was lower among patients aged ≥50 than among those aged <50 (−7.1 percentage points; p=1.9×10^-4^; q=7.6×10^-4^), whereas P3 was higher (+5.1 percentage points; p=0.0011; q=0.002). P2 and P4 showed no significant difference (Figure 1D). Therefore, the binary comparison compressed some profiles and did not define a clean molecular threshold. Resolving age by decade uncovered the structure that was compressed by the age 50 threshold (Figure 1E; Supplementary Table 5). P1 followed a significant inverted-U trajectory, peaking at 40-49 years before declining (quadratic p=4.2×10^-6^; q=8.5×10^-6^). P4 showed a U-shaped trajectory, with higher prevalence at the youngest and oldest age extremes (quadratic p=8.4×10^-15^; q=6.7×10^-14^). P3 increased linearly with age (p=3.2×10^-5^; q=5.1×10^-5^), whereas P2 decreased with age (unadjusted p=0.067; q=0.09). P2 behaviour was study-sensitive and became significant after adjustment for study, sex and tumour site (p/q=0.001). Across the seven-decade strata, mean pairwise profile separation peaked at 40-49 years (19.1 percentage points) and declined to 2.7 percentage points at 80+ years (post-peak r=-0.97, p=0.005; Figure 1F). Thus, the greatest divergence among the profiles occurred within the conventional early-onset interval, rather than at the age-50 threshold itself.

Leave-one-study-out analyses kept the P1/P4 quadratic and P3 linear age trends in all nine study omissions, whereas the P2 linear trend was maintained in four of nine (Supplementary Figure 3A). In patient-level models adjusted for study, sex and tumour site, the selected terms were significant for all profiles (P1 quadratic q=5.4×10^-4^; P2 linear q=0.001; P3 linear q=2.3×10^-4^; P4 quadratic q=2.8×10^-6^; Supplementary Figure 3B; Supplementary Table 6). Two-stage random-effects analyses confirmed the age-related trajectories for P1, P2 and P3, whereas the magnitude of the P4 quadratic pattern showed greater cross-study heterogeneity; anatomical enrichment remained highly consistent across cohorts (Supplementary Figures 4 and 5; Supplementary Table 7).

**Figure 2.**
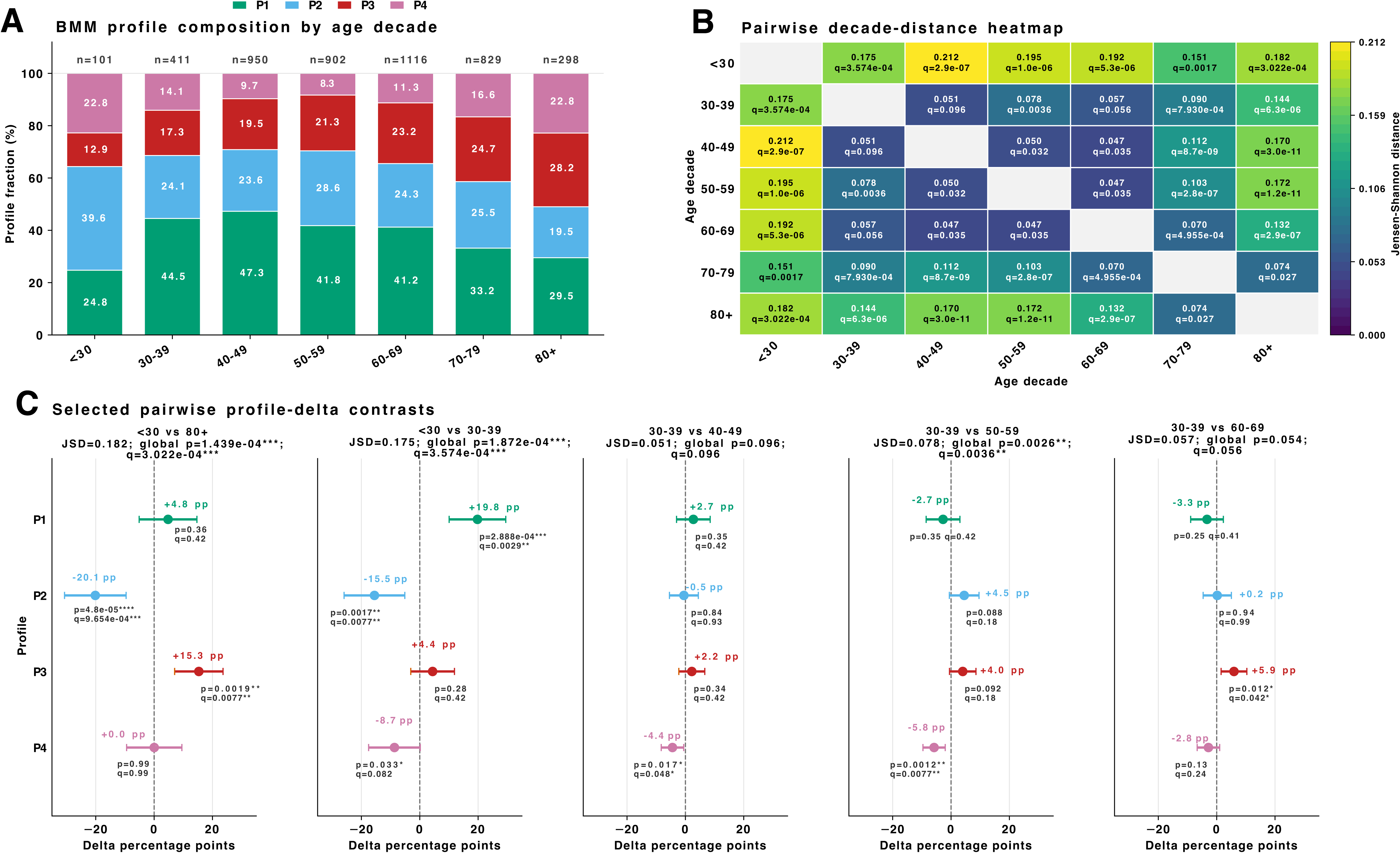
Age-decade similarity of BMM profile composition. A. Stacked P1-P4 profile composition across age decades. B. Pairwise Jensen-Shannon distances between age-decade composition vectors, with global 2 × 4 composition-test p and q values. C. Selected profile-wise contrasts testing age-extreme similarity, within-younger-onset heterogeneity and pre-versus post-50 comparisons.

**Figure 3.**
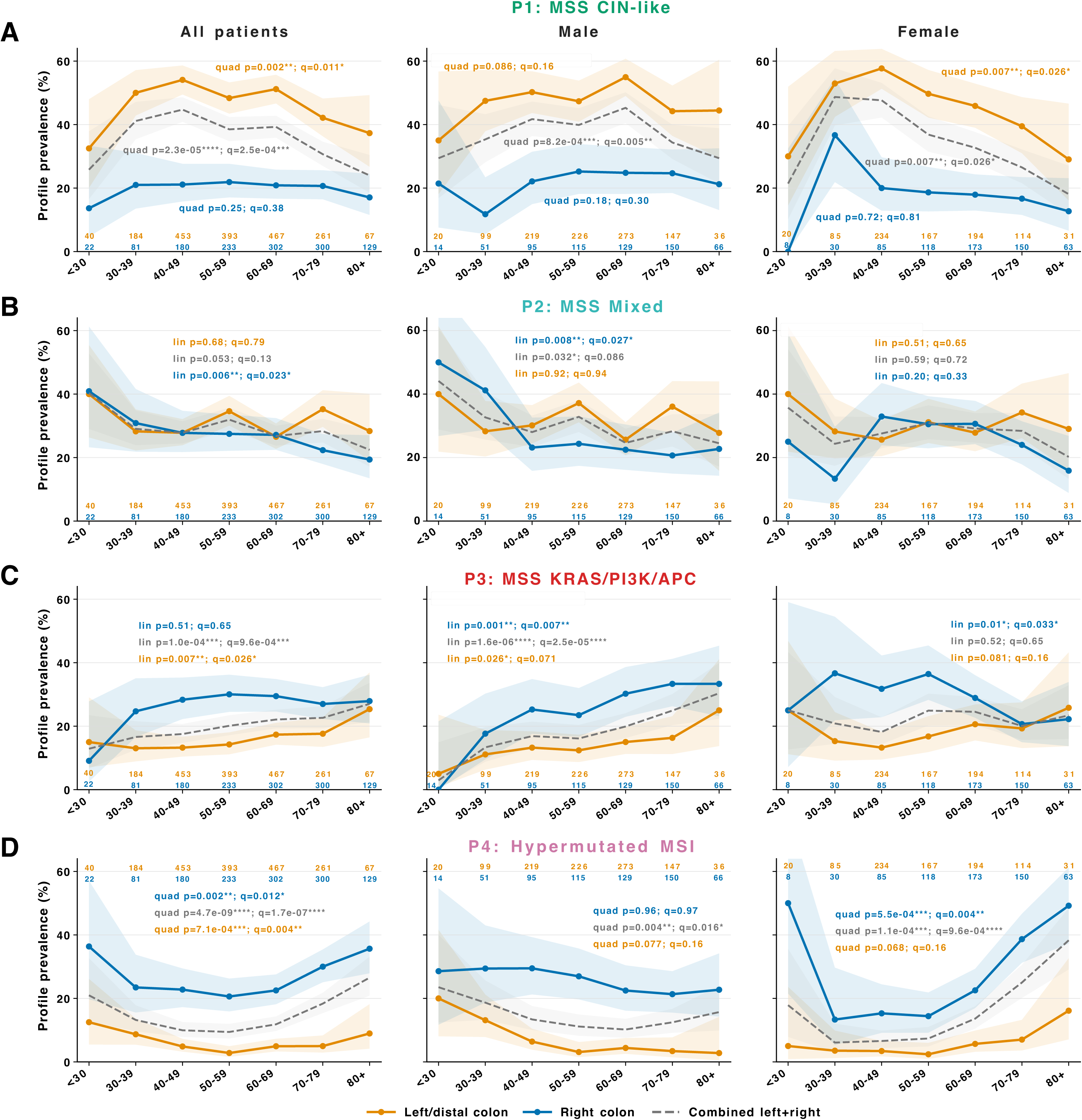
Anatomy and sex reshape profile trajectories across age decades. Rows show P1-P4; columns show all patients, males and females. Lines show profile prevalence by age decade within left/distal colon, right/proximal colon and the combined left + right population; ribbons are 95% Wilson confidence intervals. Rectal and unknown-site tumours are excluded. Sample size is indicated for each age group. P and q values for each quadratic/linear trend are indicated.

**Figure 4.**
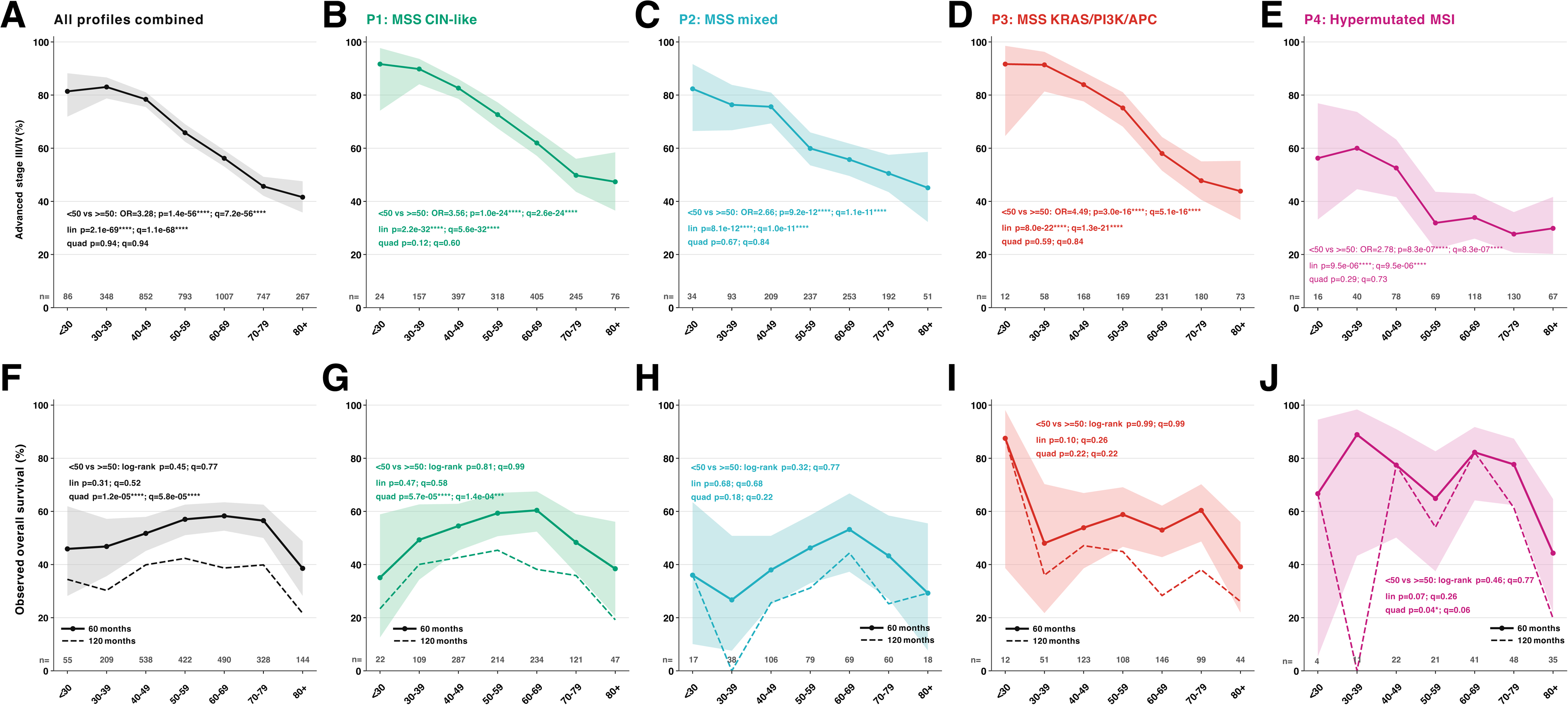
Profile-specific stage and survival patterns across age decades. A-E. Proportion of stage III/IV disease among stage-evaluable patients for all profiles combined and within P1-P4. F-J. Observed overall survival estimated by Kaplan-Meier for the same groups. Solid lines and 95% confidence ribbons show the primary 60-month estimates; dashed lines show the corresponding 120-month estimates as a longer-horizon sensitivity analysis. Lines are black/grey for all profiles combined and profile-coloured for P1-P4. Numbers immediately above the x-axis indicate the evaluable sample size within each age decade. Insets show the 60-month <50 versus ≥50 log-rank comparison and the linear and quadratic Cox age-trend statistics, with Benjamini-Hochberg-adjusted q values across the five displayed groups within each test family.

**Figure 5.**
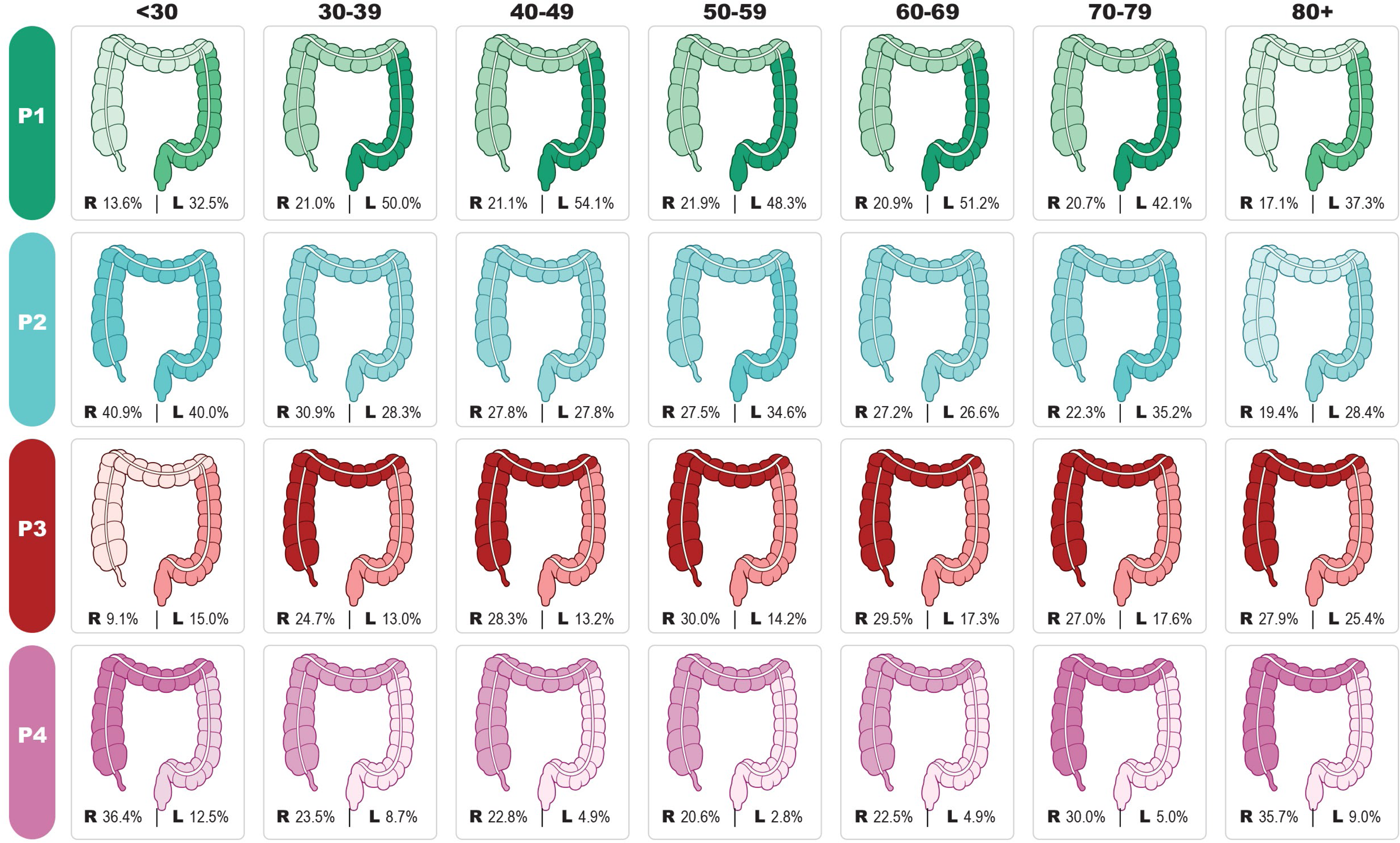
Profile-resolved anatomical prevalence across age decades. Each panel shows one BMM profile within one age decade. Viewer left represents right/proximal colon and viewer right represents left/distal colon. Colour intensity reflects profile prevalence within each side-specific stratum. Rectum is excluded. The schematic maps side-specific prevalence onto an anatomical template and should not be interpreted as segment-level sampling.

Because the nine retained cohorts combined whole-exome sequencing (WES) and targeted-panel studies, we also assessed platform sensitivity. All trends were compatible within both platform strata, whereas P2 and the magnitude of the P4 trajectory were more platform-sensitive (Supplementary Figure 6). Refitting the K=4 model after excluding FGA-high preserved 96.4% of patient assignments and reproduced the molecular centroids and age trajectories (Supplementary Figure 7), indicating that the principal architecture was not dependent on differential FGA availability.

To validate whether these conclusions depended on the BMM conditional-independence assumption or on decade binning, we fitted a four-class mixture of latent trait analysers (MLTA) and continuous-age generalised additive mixed models (GAMMs) after profile assignment (Supplementary Figure 8). MLTA reproduced the molecular centroids (overall r=0.995; mean absolute error=1.1 percentage points), matched 91.2% of patient assignments (adjusted Rand index=0.78) and preserved decade-resolved prevalence trajectories (overall r=0.953) (Supplementary Figure 8A-D). The age-50 split retained only 21.4% of decade-resolved deviance under MLTA, compared with 17.8% under BMM. Study-random-intercept GAMMs over ages 25-85 reproduced non-linear P1/P4 trajectories and opposing linear P2/P3 trends (Supplementary Figure 8E). Thus, the profile structure and non-binary age redistribution were not specific to BMM or categorical age coding.

To examine whether these profile-level trajectories were also evident in the underlying molecular features, we pooled all profiles and measured feature prevalence directly within each age decade (Supplementary Figure 9). TMB-high prevalence followed a marked U-shaped trajectory (4,607 evaluable patients; 781 events; quadratic P=1.1×10^-9^, q=1.5×10^-9^). Among 4,160 MSI-evaluable patients, MSI-H/dMMR was similarly U-shaped (452 events; quadratic P=1.8×10^-12^, q=3.7×10^-12^); by definition, its complement, MSS/pMMR/MSI-L, followed the reciprocal inverted-U pattern (quadratic P=1.8×10^-12^, q=3.7×10^-12^). By contrast, although FGA-high prevalence did show a quadratic trajectory, the trend did not reach statistical significance after correction (2,379 evaluable patients; 1,191 events; quadratic p/q=0.065). Thus, the non-linear age structure was directly visible along the hypermutation and mismatch-repair axis, whereas the copy-number axis was comparatively stable when profiles were combined.

### Distant age groups can be more compositionally similar than closer groups

We next compared the complete P1-P4 composition of all 21 pairwise decade comparisons using Jensen-Shannon distance and global 2 × 4 composition tests (Figure 2A-B). Profile composition followed neither the age-50 split nor a chronological gradient. For instance, CRC diagnosed at 30-39 years differed from 50-59 years (JSD=0.078; p=0.00259; global q=0.0036), but not clearly from 40-49 (JSD=0.051; p=0.0964; q=0.096) or 60-69 years (JSD=0.057; p=0.0535; q=0.056) (Figure 2C). Likewise, the 40-49 decade remained compositionally close to adjacent and later post-50 decades. Conversely, age <30 CRC was distinct from adjacent younger-onset decades but partially converged with 80+ diagnosed disease through similar P1/P4 prevalence while differing in P2/P3 balance. Hence, neither chronological proximity nor location on the same side of the age-50 threshold reliably predicted profile composition.

Notably, P1/P4 prevalence convergence of the youngest and oldest decades did not imply identical intra-profile biology. Within P1, the APC/MSS backbone was preserved, but broader age-extreme comparisons differed in *TP53* prevalence and sidedness (Supplementary Figure 10). Within P4, younger tumours were more APC/KRAS/MSI-high-rich, whereas older tumours were more BRAF/RNF43-rich and right-sided (Supplementary Figure 11).

Despite these variations at the age extremes, the core molecular backbones defining each profile remained robust across the lifespan. Overall, within-profile feature prevalence was largely concordant across decades and across the age-50 threshold (Supplementary Figure 12A, B). Yet the age-50 threshold comparison retained only 30% and 33% of the full decade-resolved prevalence ranges for P1 and P3, respectively, and only 3-8% for P2 and P4 (Supplementary Figure 12C). Overall, the <50/≥50 split accounted for just 17.8% of the total decade-resolved profile-composition deviance, leaving 82.2% as additional structure within the pooled groups (likelihood-ratio χ² (15) =114.0, p=2.8×10^-17^; Supplementary Figure 12D). The age-50 threshold, therefore, preserved the broad identity of the molecular states, but it poorly represented how often those states occurred across the life course.

### Anatomy and sex reshape profile trajectories

We next examined how anatomy and sex structured this age-dependent redistribution. Stratification by tumour side and sex showed that P1 remained strongly left-enriched, with a quadratic age trajectory supported in left-sided tumours overall and among females, but not males (Figure 3A). The P2 decline was concentrated in right-sided tumours and was most evident among males (Figure 3B). P3 remained relatively right-enriched and increased with age most consistently among males, particularly in the right colon, whereas female trajectories were more heterogeneous across sides (Figure 3C). P4 remained strongly right-enriched, with its U-shaped age trajectory most pronounced among females, especially in right-sided tumours (Figure 3D). Thus, anatomy and sex were associated not only with differences in average profile prevalence, but also with the shape and amplitude of their age trajectories.

### Advanced-stage presentation and survival follow distinct age patterns across molecular profiles

We next asked whether clinical presentation and survival followed the same age structure as molecular profile composition. Advanced stage III/IV at diagnosis and observed overall survival were analysed across age decades in all patients and within each BMM profile (Figure 4). Survival was evaluated primarily at 60 months, with 120-month estimates included as a longer-horizon sensitivity analysis. This profile-specific framework tested whether the clinical features associated with younger diagnosis persisted after stratification by molecular state.

Advanced-stage presentation decreased linearly with age in the full cohort (p=2.1×10^-69^; q=1.1×10^-68^). Patients aged <50 had higher odds of stage III/IV disease than those aged ≥50 (79.9% versus 54.7%; OR=3.28; q=7.2×10^-56^), and this difference persisted within every profile: +23.6 percentage points in P1 (OR=3.56; q=2.6×10^-24^), +21.5 in P2 (OR=2.66; q=1.1×10^-11^), +28.1 in P3 (OR=4.49; q=5.1×10^-16^) and +24.5 for P4 (OR=2.78; q=8.3×10^-7^). Linear age terms were significant for all four profiles (Figure 4A-E). Thus, advanced-stage presentation among younger patients was consistently observed across molecular profiles.

Observed survival followed a different pattern (Figure 4F-J). At 60 months, no <50 versus ≥50 survival contrast was detected in the full cohort or within any profile (FDR-adjusted q=0.77-0.99). Quadratic Cox age terms were significant in all patients (q=5.8×10) and P1 (q=1.4×10), but not P2 or P3; P4 showed nominal curvature that did not remain significant after FDR correction (p=0.038; q=0.063). P1 contributed 47.3% of the 2,186 survival-evaluable patients and 44.1% of deaths by 60 months (310/703). P2 showed the poorest and P4 the most favourable observed survival at 60 months. The 120-month sensitivity analysis preserved this qualitative profile ordering and strengthened the P4 quadratic signal (q=0.010). Advanced presentation and observed survival therefore represented distinct age-associated clinical axes: the former was consistently young-enriched within every profile, whereas the latter was profile-weighted and non-linear rather than defined by an age-50 contrast. The unadjusted profile-level Kaplan-Meier benchmark showed the same P2-poor/P4-favourable ordering (Supplementary Figure 13).

### Profile-resolved anatomical maps summarise the left-right profile landscape across decades

To visualise how age and anatomy jointly structured profile prevalence, we mapped the prevalence within left/distal and right/proximal tumours onto the corresponding sides of a schematic colon for each profile and age decade, excluding rectal and unknown-site tumours (Figure 5). P1 remained left/distal-dominant, with its highest left-sided prevalence from 30-39 to 60-69 years. P4 remained right/proximal-dominant and was most prevalent at the youngest and oldest extremes. P3 increased with age and was consistently more right-shifted than P1/P2, whereas P2 was highest in the youngest decade and lower thereafter. The maps recapitulate the same result regionally: molecular states are shared across age, but their anatomical representation changes across the life course.

### Rectal cancers combine a P1-high backbone with distinct composition and age trajectories

We analysed rectal cancers separately because they are anatomically distal but clinically and epidemiologically distinct from colon cancers (Supplementary Figure 14). Their dominant P1 fraction was similar to left/distal colon (50.7% versus 49.0%), but the remaining composition differed: rectal cancers were P2-depleted and P3-enriched relative to left/distal colon, while remaining P4-low relative to right colon (Supplementary Figure 14B,C). Rectum, therefore, shared a conventional P1-high backbone with left/distal colon without being compositionally equivalent to it.

Rectal profile prevalence also varied across age decades (Supplementary Figure 14C). P1 rose from the youngest group to a midlife plateau before declining (p=0.0237; q=0.0632). P3 increased nominally with age (linear p=0.0385; q=0.0714), whereas P2 remained comparatively stable (linear p=0.875; q=0.993). By contrast, P4 followed a significant lower-amplitude U-shaped trajectory (quadratic p=0.00543; q=0.0217). As in the primary colon analysis, the age-50 split captured only part of the rectal age structure, retaining 24.9% of decade-resolved profile-composition deviance and leaving 75.1% within the pooled <50 and ≥50 groups. Excluding the MSK Rectal 2022 cohort preserved the P1-high, P2-low and P3-enriched rectal composition (Supplementary Figure 15). Rectal cancer thus provided a distinct distal boundary case in which anatomy and age jointly redistributed the same molecular profiles.

### Non-overlapping MSK-IMPACT validation reproduces shared profiles and decade-resolved prevalence redistribution

Finally, to test portability beyond the discovery patients, we analysed the MSK-IMPACT 50K cohort^14^. Of 4,993 patients with a colorectal tumour record, 3,575 had a primary CRC specimen. We removed 1,340 patients whose MSK patient identifiers matched any discovery MSK cohort (crc_eo_2020, crc_msk_2017 or rectal_msk_2022) and retained one primary specimen per patient, yielding 2,235 non-overlapping validation patients; age at diagnosis was available for 2,134, including 502 diagnosed before age 50 (Supplementary Figure 16A). Using profile-specific Bernoulli probabilities fixed from the discovery assignments, we projected the four BMM profiles into the validation patients based on 30 shared molecular features (ACVR2A was absent), without using age, sex or tumour site. Profile-specific molecular-feature prevalence patterns remained strongly concordant between discovery and validation (r=0.959-0.998; overall r=0.984; Supplementary Figure 16B).

The decade-resolved prevalence trajectories were also strongly concordant with the discovery cohort (within-profile standardised r=0.904, p=4.38×10^−11^), with profile-specific correlations of 0.967 for P1, 0.854 for P2, 0.836 for P3 and 0.959 for P4 (Supplementary Figure 16C). In models adjusted for sex, tumour site and IMPACT panel, P1/P4 remained quadratic (q=0.002 and 1.4×10−4, respectively), P3 retained a linear association (q=0.033), and P2 retained the negative linear trend (q=0.067; Supplementary Figure 16D). The conventional <50/≥50 split retained only 6.2% of decade-resolved profile-composition deviance, lower than the 17.8% in the discovery cohort. Scanning all possible integer age cut-offs confirmed that binary representation remained incomplete even at the best-performing thresholds, which occurred at age 70 in discovery (51.3% of decade-resolved deviance) and age 75 in validation (49.4%; Supplementary Figure 16E).

Importantly, individual pairwise contrasts were not anchored to age 50. In the validation cohort, CRC at 30-39 years did not differ from either CRC at 50-59 years (q=0.258) or CRC at 60-69 years (q=0.478), whereas <30 and 80+ disease remained globally distinct (q=0.029) despite partial convergence in P1/P4 prevalence. The direction and overall shape of the projected trajectories were preserved after excluding patients with another recorded cancer, restricting to assignments with maximum posterior probability ≥0.80 and calculating FGA from autosomes only. Thus, the non-overlapping validation cohort reproduced the age-dependent redistribution of shared profiles.

## Discussion

While molecular subtyping frameworks have resolved distinct biological states in colorectal cancer (CRC)^1,2,4,15^, epidemiological studies of rising early-onset incidence commonly use age 50 to distinguish early- from average-onset disease^8–12,16–22^. Here, by resolving molecular-profile prevalence at diagnosis across life decades, we show that this epidemiologically useful threshold captures only a limited part of the age-associated structure: shared profiles retain their molecular identities but follow distinct linear and non-linear prevalence trajectories. Our findings therefore complement the early-versus average-onset framework by providing a profile-resolved view of CRC biology across the life course.

The four profiles, identified without age, sex or tumour site, mapped onto established CRC axes and remained recognisable across age^2,4^. Their recovery across all retained cohorts and persistence in leave-one-study-out, covariate-adjusted, platform and FGA sensitivity analyses argue against dependence on a single cohort or incompletely measured feature. An MLTA reproduced their molecular identities and age-decade prevalence, continuous-age GAMMs recovered the prespecified P1/P4 curvature and P2/P3 linear trends, and both molecular patterns and age trajectories were reproduced in 2,235 non-overlapping MSK-IMPACT patients^14^. In the corrected validation cohort, three of four prespecified age terms retained FDR-adjusted support and the age-50 split retained only 6.2% of decade-resolved deviance. Together, these analyses show that profile recovery and age structure were robust to modelling assumptions, age parameterisation and dataset.

Binary age representation was incomplete regardless of the selected cut-off. The age-50 threshold retained only 17.8% and 6.2% of decade-resolved compositional deviance in discovery and validation, respectively, while even the best-performing thresholds retained only 51.3% and 49.4%. Thus, profile identities remained stable while their prevalence changed across decades, so any single threshold compressed much of the underlying structure. Although age 50 remains useful for screening, epidemiology and communication^17–19,23^, treating it as a molecular threshold obscures variation within both early- and average-onset disease.

Age thresholds serve different purposes in epidemiology and molecular classification. Population-level incidence trends and screening benefit-risk appropriately inform screening starting ages, whereas our analysis characterises molecular-profile prevalence among CRCs diagnosed at different ages. Lowering the recommended starting age to 45 years is an important public-health response but does not create a second molecular threshold^23^. Associations with age at diagnosis may jointly reflect biological ageing, birth cohort, calendar period, screening, diagnostic pathways and exposures, which cross-sectional data cannot disentangle.

More than half of patients diagnosed before age 40 came from a cohort diagnosed in 2014-2019, placing its 30-39-year-old patients mainly in birth cohorts from the mid-1970s through the 1980s. Adults currently around age 30 were born in the mid-1990s, while many people currently under 30 were born in the 2000s; these trajectories should therefore be reassessed in newer birth cohorts. Rather than motivating additional molecular age thresholds, these findings support treating age as a continuous or decade-resolved dimension of CRC molecular epidemiology. Modelling CRC biology across age in this way may better resolve age-associated molecular heterogeneity and biologically meaningful subgroups without imposing arbitrary cut-offs.

Although younger patients with CRC more often present with advanced-stage disease and other high-risk clinicopathological features^9–12,17–20,24,25^, this cannot be explained solely by enrichment for one adverse molecular profile. Advanced-stage presentation declined with age within P1, P2, P3 and P4, indicating that younger presentation is a clinical axis superimposed on the molecular-profile architecture. Observed 60-month survival followed a distinct, profile-weighted and non-linear pattern, without a robust age-50 contrast; the 120-month sensitivity analysis preserved the same qualitative profile ordering. Advanced-stage presentation may reflect delays both before and after first clinical contact, including delayed symptom appraisal and lower clinical suspicion of CRC in younger adults^24^, which may contribute to advanced presentation. However, neither our cross-sectional data nor stage at diagnosis directly measures tumour growth kinetics or provides a proxy for a shorter tumour ‘age’ in younger patients^20,22^. Stage, molecular composition and survival are therefore related but non-interchangeable. This argues against interpreting early-onset CRC as intrinsically more aggressive on the basis of age alone: outcomes depend on tumour biology, stage, anatomical location, timely diagnosis and access to optimal treatment^25,26^.

Pairwise comparisons further show why no single cross-threshold contrast defines the biology. In the discovery cohort, profile composition at 30-39 years differed from that at 50-59 years but not clearly from that at 60-69 years; in the MSK validation cohort, 30-39-year CRC differed from neither post-50 decade after multiplicity correction. Thus, the reproducible feature was the redistribution trajectory, not a fixed cross-threshold contrast. At the age extremes, <30 and 80+ disease remained globally distinct in both cohorts but partially converged through similar P1/P4 prevalence. This convergence did not imply molecular equivalence: P1 retained its APC/MSS backbone but differed in TP53 prevalence and sidedness, whereas P4 shifted from a younger APC/KRAS/somatic-MMR-alteration-rich context to an older BRAF/RNF43-rich, right-sided context^27–35^. These findings are compatible with distinct biological routes to shared profile states.

Although the profile framework is concordant with existing CRC biology and aligned with the genomic and mismatch repair-burden components of CMS, it is not a CMS substitute. P4 was closest to CMS1, P1 to CMS2 and P3 to CMS3, consistent with their hypermutated/MSI-high, conventional/CIN-like and KRAS/PI3K/APC-rich backbones. P2 showed only weak or partial proximity to CMS4, as expected because CMS4 is defined largely by mesenchymal, stromal and TGF-β-associated transcriptomic biology rather than a distinctive somatic-driver configuration^2,4,15^. These similarities indicate that the BMM recovered established genomic backbones rather than transcriptional CMS identities, enabling their consistent interrogation across age, anatomy and sex. A previous study described similar U-shaped CMS1 and inverted-U CMS2 patterns across age at diagnosis^9^, although it did not formally test these trends.

Anatomy was not merely a validation feature but a coordinate of the age trajectories. P1 remained concentrated in left/distal disease, whereas P4 remained concentrated in right/proximal disease. P3 occupied a progressively older and more right-shifted state, while P2 remained intermediate. Sex further amplified tendencies present in all patients, including the non-linear prevalence of left-sided P1 and right-sided P4 among females, reinforcing the relevance of biological sex to cancer risk, progression and treatment response^5–7,36–39^. This spatial segregation aligns with established biological differences between the right and left colon, which derive from distinct embryonic origins and maintain different physiological, molecular and microbial environments^40,41^.

Rectal cancer provided an informative anatomical boundary case. Its dominant P1 component was left-like, but its non-P1 compartment was not equivalent to left/distal colon: rectum was P2-depleted and P3-enriched while remaining P4-low relative to right colon. This pattern persisted after excluding the largest rectal-specific cohort. Keeping rectum separate from the primary left-right contrast was therefore anatomically appropriate and prevented a distinct distal composition from being absorbed into the left-colon category^5,42^.

Although the identified profiles are not directly treatment-defining, their coexistence across age groups shows that early- or average-onset status is not a surrogate for an individual tumour’s molecular state. Current guidelines base therapeutic selection on established molecular biomarkers rather than age alone^43^, consistent with evidence that chemotherapy response and survival among patients with advanced microsatellite-stable CRC can be comparable across age groups^26^. Future biomarker-driven trials should therefore model age and molecular-profile composition jointly rather than use the <50/≥50 distinction as a proxy for tumour biology. Whether these profiles explain or predict treatment-response heterogeneity beyond established biomarkers requires prospective testing. Their clinical utility beyond current biomarkers, staging and risk classifications also remains to be established.

The main limitations of this study are those of retrospective public cohorts: heterogeneous sequencing platforms and study-specific age and site distributions, incomplete clinical annotation and missing germline, methylation, mutational-signature, microbiome, exposome, screening, recurrence and treatment data. Patient-level diagnosis year was unavailable for most cohorts, preventing formal separation of age, calendar-period and birth-cohort effects. Contemporary young adults born in the mid-1990s or 2000s were underrepresented or represented only within the youngest age group, limiting extrapolation to current cohorts.

Race and ethnicity annotations were cohort-dependent and insufficient for causal disparities analyses. Observed overall survival reflected all-cause mortality and was not adjusted for stage or treatment; treatment exposure was not harmonised across cohorts, so the contributions of stage and specific therapeutic modalities could not be disentangled. Platform-stratified analyses, the no-FGA refit and non-overlapping MSK-IMPACT validation reduce concern that the principal profiles were artefacts of cohort composition, sequencing strategy or FGA availability. However, the validation cohort was derived from the same institution and assay ecosystem as three discovery cohorts, despite complete patient-level overlap exclusion. We also evaluated AACR Project GENIE as a potential external validation resource; however, the CRC BPC cohort was dominated by MSK and DFCI, both represented in the discovery dataset, whereas the broader GENIE Registry records age at sequencing rather than age at initial diagnosis, precluding a fully institutionally independent, diagnosis-anchored replication.

In conclusion, established CRC molecular programmes persist across the life course while their prevalence, anatomical context and clinical presentation follow profile-specific linear and non-linear age trajectories. Age 50 remains useful for screening, epidemiology and communication but is not a molecular threshold. The reproduction of profile identities and prevalence trajectories in non-overlapping validation patients supports modelling age as a trajectory alongside anatomy, sex and study context, rather than treating CRCs on either side of age 50 as two uniform diseases.

## Methods

### Study design and public-cohort assembly

This was a retrospective, harmonised, patient-level analysis of public CRC sequencing cohorts available through cBioPortal. We screened 28 CRC-relevant studies and retained nine after applying eligibility, age-availability, primary-tumour, duplication and harmonisation criteria (Supplementary Table 1). The final discovery dataset included 4,609 unique patients with primary CRC. Study-prefixed patient and sample identifiers were used to prevent identifier collisions; duplicate sample rows were removed, one eligible primary molecular record was retained per patient and TCGA-derived records were collapsed to a single PanCancer Atlas release. Disease-context-enriched, treatment-specific and non-primary CRC cohorts were excluded.

The harmonised table included study identifier, sequencing-platform group, age at diagnosis, sex, race and ethnicity when available, tumour site, stage, MSI/MMR status, TMB, FGA, survival annotation and gene-level binary alteration calls. Tumour site was classified as right/proximal colon, left/distal colon, rectum or unknown. Caecum, ascending colon, hepatic flexure and transverse colon were assigned to the right/proximal group; splenic flexure, descending colon and sigmoid colon were assigned to the left/distal group. Rectal and unknown-site tumours were excluded from primary left-right analyses, and rectal cancers were analysed separately.

All records were deidentified and publicly available. Ethical oversight and consent for the original data collections were reported by the contributing studies. No statistical method was used to predetermine sample size; all eligible patients were included. Randomisation and blinding were not applicable.

### Molecular-feature harmonisation and BMM input matrix

The BMM input matrix comprised 31 binary molecular features: MSI-H/dMMR, FGA-high, TMB-high and qualifying non-silent somatic mutation indicators for APC, TP53, KRAS, NRAS, BRAF, RNF43, PIK3CA, PTEN, SMAD4, SMAD2, TGFBR2, ACVR2A, FBXW7, SOX9, CTNNB1, TCF7L2, ARID1A, KMT2D, ATM, ERBB2, GNAS, POLE, POLD1, MLH1, MSH2, MSH6, PMS2 and EPCAM. These features spanned repair state, genomic burden, core CRC drivers and pathway alterations while remaining interpretable across WES/WXS and targeted-panel cohorts. Age, sex and tumour site were excluded from model fitting and introduced only in downstream analyses.

TMB-high was defined as TMB ≥10 mutations/Mb. FGA-high was defined within each study as FGA greater than or equal to the study-specific median among FGA-evaluable patients after cohort-level age filtering. MSI-H/dMMR was derived from harmonised study-provided MSI/MMR fields; MSS/pMMR was defined as its complement among evaluable tumours. Missing MSI/MMR and FGA values were retained as missing and omitted from the corresponding Bernoulli likelihood contributions; no missing category was created and no molecular feature was imputed. TMB and gene-mutation indicators were complete in the discovery dataset.

### Bernoulli mixture modelling and profile assignment

Bernoulli mixture models were fitted to the binary molecular-feature matrix for K=2-5. Conditional on latent profile membership, each observed feature was modelled as a Bernoulli variable with a profile-specific probability, and patients were assigned to the component with the highest posterior responsibility. We inspected log-likelihood, AIC, BIC, positive-form ICL, entropy, mean maximum posterior probability, the proportion of patients with posterior probability ≥0.80 and cluster sizes (Supplementary Figure 1; Supplementary Table 3). K=4 was retained as the principal broad resolution because it produced four interpretable and adequately sized profiles for seven-decade, anatomy-, sex- and endpoint-resolved analyses. K=5 improved likelihood-based criteria but generated a 66-patient component, so it was not used as the primary population-level resolution.

### CMS-BMM profile proximity analysis

CMS-compatible expression data were not uniformly available across the retained cohorts, precluding harmonised patient-level CMS assignment. We therefore compared profile-level molecular vectors rather than classifying individual tumours. CMS feature percentages were taken from Supplementary Table 5 in Guinney et al.^4^, and BMM percentages from the K=4 feature-prevalence matrix. Anatomical side was omitted from the feature vectors; no patients were excluded on the basis of tumour location. The eight shared features were MSI-H/dMMR, TMB-high/hypermutation, FGA-high/SCNA-high, KRAS, BRAF, PIK3CA, TP53 and APC. Pearson and Spearman correlations, feature-wise z-scored Euclidean distance and z-scored cosine similarity were calculated for all 16 BMM-CMS pairs; Pearson p values were adjusted using Benjamini-Hochberg FDR.

### Age coding and trend tests

Age at diagnosis was represented either by seven groups (<30, 30-39, 40-49, 50-59, 60-69, 70-79 and 80+) or as continuous age centred and scaled in 10-year units. For Figure 1E, one-versus-rest logistic models tested linear and quadratic age terms for every profile, with Benjamini-Hochberg correction across the eight unadjusted profile-term tests. The trajectory-matched terms displayed in Figure 1E were quadratic for P1/P4 and linear for P2/P3; both terms for all profiles are reported in Supplementary Table 5. The same four terms were specified before analysis of the validation cohort. Sensitivity models additionally adjusted for study, sex and tumour site.

### Profile prevalence and age-decade composition

For each profile, age group and analysis stratum, prevalence was calculated as the number assigned to that profile divided by the total number in the stratum. Wilson 95% confidence intervals were used for plotted proportions. Profile-specific <50 versus ≥50 contrasts used Fisher exact tests with Benjamini-Hochberg correction across the four profiles. For Figure 2, each decade was represented by its four-profile composition vector. Jensen-Shannon distance using natural logarithms and total variation distance were calculated for all 21 pairwise decade comparisons, and global composition differences were tested using 2 × 4 chi-square tests with Benjamini-Hochberg correction across the 21 pairwise tests. Selected profile-wise contrasts used two-proportion tests with 95% confidence intervals and FDR correction within the displayed contrast family; the younger decade was the reference, so positive values indicate greater prevalence in the comparison decade.

For Figure 1B, right/proximal versus left/distal location was regressed on an ordered profile score (P1=1, P2=2, P3=3 and P4=4) in a logistic model; rectal and unknown-site tumours were excluded. For Figure 1F, the six absolute pairwise differences among P1-P4 prevalence were calculated within each age decade and averaged. The post-peak association from 40-49 years through 80+ was summarised by Pearson correlation.

### Direct molecular-feature age trajectories

To determine whether profile-level trajectories were mirrored by the underlying state and burden features, TMB-high, FGA-high, MSI-H/dMMR and MSS/pMMR prevalence were analysed across age decades (Supplementary Figure 9). Available-case denominators were used. Logistic models included linear age in 10-year units, and nested likelihood-ratio tests compared linear with linear-plus-quadratic age models. Benjamini-Hochberg correction was applied across the four feature-specific non-linearity tests.

### Age-extreme molecular-context analyses

Within P1 and P4, selected molecular and anatomical features were compared between <30 and 80+ years, with <40 versus ≥70 years used as a broader sensitivity contrast (Supplementary Figures 10 and 11). Effects were expressed as older-minus-younger prevalence differences with Newcombe 95% confidence intervals. Fisher exact p values were adjusted within each displayed profile-comparison family using Benjamini-Hochberg FDR. FGA-high used available-case denominators. Somatic MMR-gene, POLE/POLD1, BRAF/RNF43 and anatomical indicators were treated as route-compatible proxies; they were not interpreted as evidence of germline status, MLH1 methylation, CIMP or specific exposure histories.

### Intra-profile molecular concordance and information retained by the age-50 threshold

To separate preservation of profile identity from redistribution of profile prevalence, intra-profile and inter-profile age effects were analysed separately (Supplementary Figure 12).

For each profile and age decade, prevalence was calculated for all 31 BMM inputs and compared with the corresponding profile-wide prevalence vector. Agreement was summarised by Lin’s concordance correlation coefficient and the mean absolute prevalence difference across features. Median values across the seven decades were reported for each profile.

The same analysis compared the 31-feature prevalence vectors of patients aged <50 and ≥50 within each profile. Because these features contributed to BMM construction, this analysis assessed internal consistency of the fitted profile architecture across age rather than independent validation of profile assignment.

For each profile, the absolute <50 versus ≥50 prevalence difference was compared with the full peak-to-trough range across seven decades. Their ratio was reported as the percentage of the observed decade-resolved range retained by the binary comparison. These range-retention values were descriptive effect-size summaries.

The total association between age decade and profile composition was quantified using likelihood-ratio deviance from the 7 × 4 decade-by-profile table. The component represented by age 50 was obtained from the 2 × 4 binary table. Additional decade-resolved structure was tested by comparing the unrestricted seven-decade model with a constrained model in which the three <50 decades shared one composition and the four ≥50 decades shared another. The binary and additional components were expressed as proportions of total decade-resolved deviance, not as variance explained.

To test whether a different binary cut-off represented the decade-resolved association more completely, the 2 × 4 likelihood-ratio deviance was recomputed for every integer threshold within the observed age range that produced non-empty lower- and upper-age groups. Retention at threshold c was calculated as 100 × G²(c) divided by the fixed 7 × 4 decade-resolved deviance. The maximum-retention threshold was treated as a descriptive optimum rather than an inferred or validated biological cut-off.

### Anatomy, sex and profile-resolved anatomical mapping

For Figure 3, profile prevalence was estimated by age decade within left/distal colon, right/proximal colon and the combined left + right population, separately for all patients, males and females. Wilson intervals were used throughout; rectal and unknown-site tumours were excluded. Within each sex-by-site stratum, one-versus-rest logistic models tested linear, quadratic and <50/≥50 age terms for each profile. Benjamini-Hochberg correction was applied across all tests displayed in Figure 3; P1/P4 quadratic and P2/P3 linear terms were used to describe the primary trajectories. Figure 5 mapped the same side-specific prevalence estimates onto a schematic colon template, with colour intensity representing prevalence within the side-specific stratum.

### Rectal cancer sensitivity analysis

Rectal cancer was treated as a separate distal compartment. Overall composition was compared with left/distal colon, combined colon and right/proximal colon using profile-wise Fisher tests and global composition tests; profile-wise p values were Benjamini-Hochberg adjusted within each four-profile comparator family. Rectal age-decade trajectories were estimated with Wilson intervals. One-versus-rest logistic models used the same trajectory-matched terms as the primary analysis (quadratic for P1/P4 and linear for P2/P3), with Benjamini-Hochberg correction across the four selected terms. The analysis was repeated after excluding the MSK Rectal 2022 cohort to assess dependence on the largest rectal-specific study (Supplementary Figures 14 and 15).

### Profile-specific stage and survival analyses

Stage annotations were mapped to major stages I-IV where possible; records without an evaluable major stage were excluded. Advanced stage was defined as stage III/IV. For Figure 4A-E, advanced-stage prevalence was estimated across age decades for all profiles combined and within P1-P4, with Wilson intervals. <50 versus ≥50 comparisons used Fisher exact tests and odds ratios, with FDR correction across the five displayed groups. Linear and quadratic age terms were tested using logistic regression with age scaled in 10-year units; p values were FDR adjusted across the five displayed groups separately for each term. For Figure 4F-J, overall-survival time and vital-status annotations followed the source-study definitions. Patients required paired survival time and event information, with death as the event. The primary analysis administratively censored follow-up at 60 months; Kaplan-Meier estimates at 60 months used Greenwood-based 95% confidence intervals. <50 versus ≥50 comparisons used log-rank tests, and linear and quadratic age terms were tested using Cox proportional-hazards models on the same 60-month-censored data, with Benjamini-Hochberg correction across the five displayed groups separately for each test family. The same baseline survival-evaluable population was re-analysed with administrative censoring at 120 months as a longer-horizon sensitivity analysis. Figure 4 displays the 60-month estimates and confidence ribbons as the primary curves and the corresponding 120-month estimates as dashed sensitivity overlays; inferential statistics displayed in Figure 4 correspond to the 60-month analysis.

Unadjusted overall survival was also compared across P1-P4 among patients with available survival time and event annotation (Supplementary Figure 13). Kaplan-Meier curves and numbers at risk were estimated by profile, with a global log-rank test and descriptive pairwise comparisons adjusted by Benjamini-Hochberg FDR. This benchmark was not adjusted for age, stage, site, study, platform or treatment.

### MLTA and generalised additive mixed-model sensitivity analyses

To test whether profile recovery depended on the BMM assumption of conditional independence among molecular features within profiles, we fitted a four-class, one-dimensional common-slope MLTA to the same 31 binary features, again excluding age, sex and tumour site. Missing MSI/MMR and FGA values were omitted from the corresponding likelihood contributions. K=4 was fixed to compare molecular states at the same resolution as the principal BMM; MLTA classes were mapped one-to-one to BMM profiles by maximum patient overlap only after fitting. Agreement was summarised using matched assignments, adjusted Rand index, profile-centroid Pearson correlation, mean absolute error and profile-by-decade prevalence correlation. The age-50 share of decade-resolved deviance was recomputed for MLTA assignments, with 95% confidence intervals from 2,000 patient-bootstrap resamples within study while preserving study sizes.

After BMM and MLTA assignments were fixed, continuous-age prevalence was modelled separately for membership in each profile versus the other three profiles using binomial GAMMs with a logit link and a Gaussian random intercept for study. Age was represented by a linear term plus four orthogonal low-rank cubic B-spline terms; the nested linear mixed model retained only the linear term. Models were estimated using a Laplace approximation around the posterior mode with weak Gaussian regularisation of fixed effects. Non-linearity was tested from the joint contribution of the four spline terms beyond linear age. Benjamini-Hochberg correction was applied across the eight BMM/MLTA non-linearity tests and, for the prespecified P2/P3 linear hypotheses, across the four method-by-profile linear-age tests. Population-level predictions set the study random intercept to zero. The displayed analysis was restricted to ages 25-85 years because age coverage was sparse at the extremes; complete-range and 30-80-year analyses, and models additionally adjusted for sex and tumour site, were used as sensitivity analyses.

### Leave-one-study-out and adjusted robustness analyses

Each retained study was omitted in turn and the selected Figure 1E terms were recomputed: P1 quadratic, P2 linear, P3 linear and P4 quadratic. Significance retention was summarised as the number of leave-one-study-out runs with q<0.05. P2 was also assessed for a quadratic alternative. Patient-level sensitivity models included the selected age term with fixed effects for study, sex and tumour site (Supplementary Figure 3; Supplementary Table 6).

### Study-level heterogeneity and random-effects sensitivity analysis

Two-stage random-effects analyses quantified study-level consistency and heterogeneity without replacing the primary pooled patient-level models. For anatomy, study-specific log odds ratios compared right/proximal with left/distal enrichment for each profile. For age, the selected profile-specific coefficient was estimated within each study using common age scaling. Estimates were combined by inverse-variance random-effects models; heterogeneity was summarised using Cochran’s Q, tau-squared (τ²), I-squared (I²), and 95% prediction intervals (Supplementary Figures 4 and 5; Supplementary Table 7).

### Sequencing-platform and FGA sensitivity analyses

For Supplementary Figure 6, the fixed principal K=4 assignments were stratified by WES/WXS versus targeted-panel profiling. Age-decade trajectories were estimated within each platform, and patient-level models tested age-by-platform interactions for the prespecified profile terms. Within-profile molecular-feature prevalence vectors were compared across platforms using concordance and mean absolute differences; feature evaluability was audited explicitly, with particular attention to FGA-high. Assignments were not refitted in this analysis.

For Supplementary Figure 7, the BMM was refitted at K=4 after excluding FGA-high, using the remaining 30 inputs and multiple initialisations. Refit components were aligned to the principal profiles by molecular-centroid similarity. Robustness was summarised using assignment retention, adjusted Rand index, normalised mutual information, centroid concordance and similarity of age-decade trajectories. Information criteria were not compared between the 31- and 30-feature models because their likelihoods were defined on different feature spaces.

### Non-overlapping MSK-IMPACT validation

We used the 2026 MSK-IMPACT 50K cBioPortal release as a patient-level validation resource^14^. Patients with a colorectal cancer record were restricted to primary specimens, and colorectal carcinoma in situ was excluded. Every MSK patient identifier present in any discovery MSK cohort (crc_eo_2020, crc_msk_2017 or rectal_msk_2022) was removed by exact patient-ID matching. When multiple eligible primary specimens were available, we selected one by prioritising FGA evaluability, passing FACETS quality control, MSI and TMB evaluability, broader IMPACT panel coverage and the earliest sample ordinal. This yielded 2,235 non-overlapping patients, including 2,134 with age at diagnosis.

Validation features followed the discovery definitions wherever possible. MSI-H was derived from study-provided “instable” versus “stable” classifications, TMB-high was defined as ≥10 mutations/Mb, and FGA was calculated as the fraction of recorded segment bases with an absolute segment mean ≥0.2. FGA-high was defined using the validation-cohort median among patients evaluable for both age and FGA. Functional somatic-mutation indicators were reconstructed for the 28 discovery genes, accounting for panel-specific missingness. ACVR2A was absent from all IMPACT panels and was marginalised rather than coded as wild type. Age, sex, tumour site and clinical outcomes were excluded from profile assignment.

For transport, profile-specific Bernoulli feature probabilities were estimated from the final discovery assignments using Jeffreys smoothing among patients evaluable for each feature, while discovery profile prevalences were used as prior probabilities; both were then frozen for validation. For each validation patient, missing features were omitted from the likelihood calculation, posterior probabilities were calculated across P1-P4 using the remaining features, and the patient was assigned to the profile with the highest posterior probability. When reapplied to the discovery cohort, this frozen projector recovered 97.9% of the original BMM assignments.

Molecular portability was quantified using Pearson and Spearman correlations between discovery and validation profile-specific feature-prevalence vectors across the 30 shared features. Profile prevalence and Wilson confidence intervals were estimated by age decade. Prespecified continuous-age terms were quadratic for P1/P4 and linear for P2/P3. For Supplementary Figure 16D, discovery models were adjusted for sex, tumour site and sequencing-platform group, whereas validation models were adjusted for sex, tumour site and IMPACT panel; Benjamini-Hochberg correction was applied across the four prespecified terms within each cohort. Trajectory concordance was calculated across the seven decades within each profile. For the pooled comparison, prevalence values were standardised separately within each profile across its seven-decade points before the 28 points were combined. The proportions of decade-resolved likelihood-ratio deviance retained by age 50 and by every integer cut-off in the observed age range were recomputed as in discovery. Sensitivity analyses excluded patients with another recorded cancer, restricted assignments to a maximum posterior probability ≥0.80 and recalculated FGA using autosomal segments only. Pairwise decade-composition tests used the same natural-logarithm Jensen-Shannon distance, chi-square tests and false-discovery-rate correction as the discovery analysis.

### Statistical analysis and reporting

All hypothesis tests were two-sided. Effect estimates are reported with 95% confidence intervals where applicable, and Benjamini-Hochberg-adjusted q values are reported for the prespecified families described above. Statistical significance was defined as p<0.05 or q<0.05, as appropriate. Analyses used available data for the variables required by each model; no clinical or molecular values were imputed. Exact analysis denominators and both nominal and adjusted p values are provided in the figure source tables and statistical output files.

### Formula compendium and reproducibility

A separate Supplementary Methods Formula Compendium provides the exact definitions and formulas used throughout the study, including Bernoulli mixture likelihoods, missing-feature likelihood contributions, posterior profile assignment, information criteria, entropy and ICL, Wilson intervals, Fisher exact and chi-square tests, Jensen-Shannon and total variation distances, profile contrasts, linear and quadratic age-trend models, Lin’s concordance correlation coefficient, mean absolute prevalence difference, age-threshold range-retention measures, likelihood-ratio deviance partitioning and threshold scanning, Kaplan-Meier estimates, log-rank tests, Cox models, leave-one-study-out retention, CMS-BMM proximity metrics, random-effects meta-analysis, Cochran’s Q, tau-squared (τ²), I-squared (I²), prediction intervals, platform interactions, assignment-similarity metrics, mixture-of-latent-trait likelihoods, cubic-spline GAMMs, study-level random effects, joint non-linearity tests and Benjamini-Hochberg FDR correction. Reproducible scripts, source tables, model assignments, statistical outputs and audit files accompany the supplementary materials.

## Data availability

All source cohorts were publicly available through cBioPortal and are identified by study ID in Supplementary Table 1. The analysis used TCGA PanCancer Atlas data and the 2026 MSK-IMPACT 50K cBioPortal release as described above. The deidentified harmonised patient-level master, study inclusion/exclusion ledger, profile assignments, figure source tables, q-value tables and validation data are supplied as Supplementary Data.

## Code availability

The complete end-to-end reproduction script, Supplementary Methods Formula Compendium, model-assignment tables, statistical outputs and audit files are supplied as Supplementary Code. Starting from the harmonised patient-level master, these materials reproduce all main and supplementary figures and analyses reported here.

---

**Supplementary Table 1.**
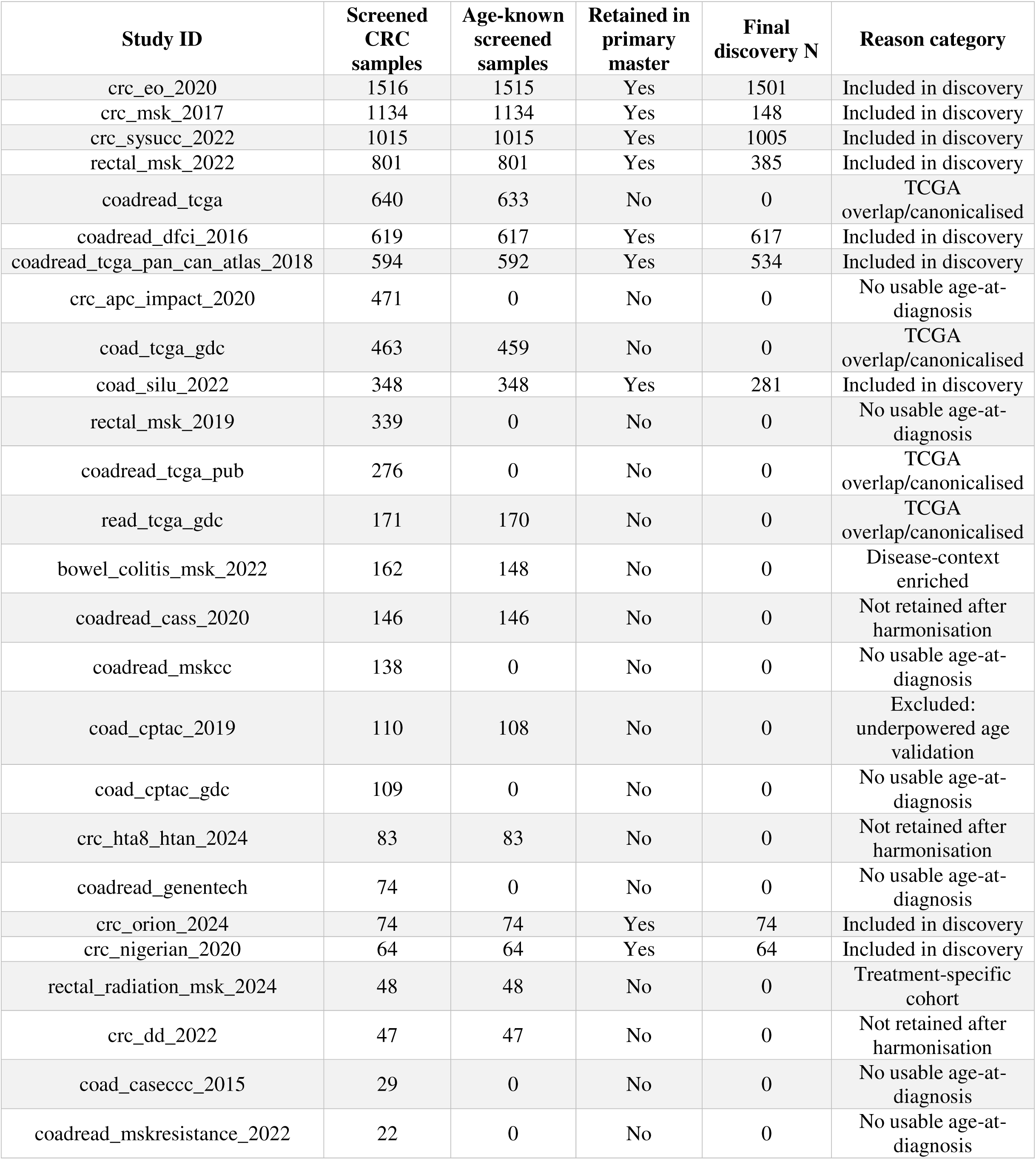
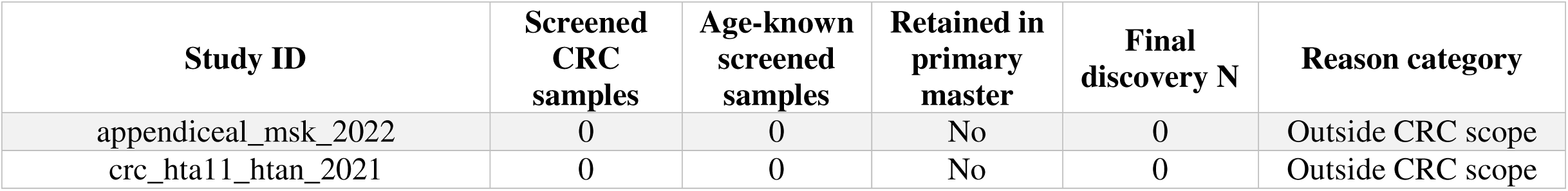
Public cBioPortal study screening, inclusion and exclusion audit.

**Supplementary Table 2.**
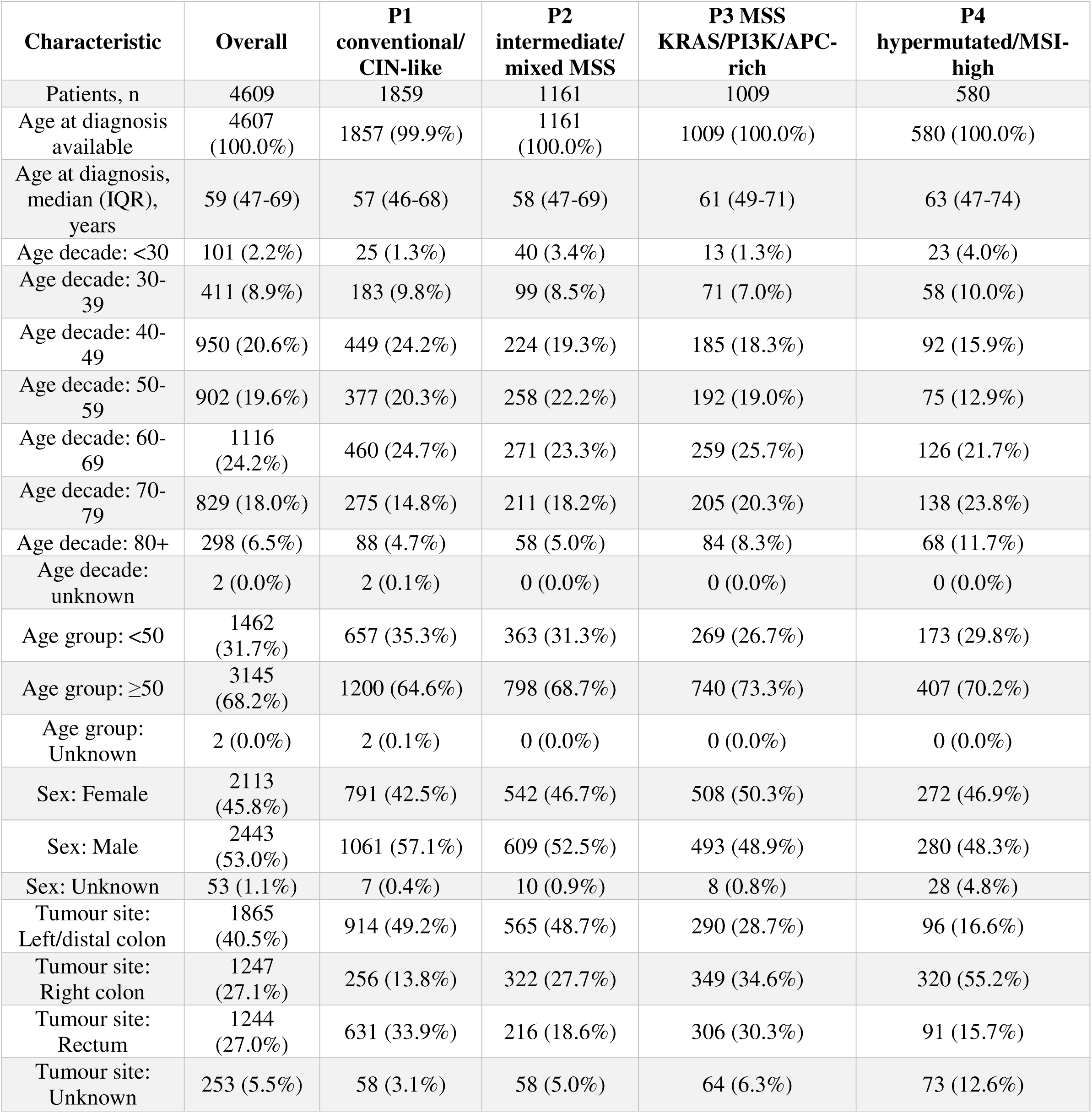
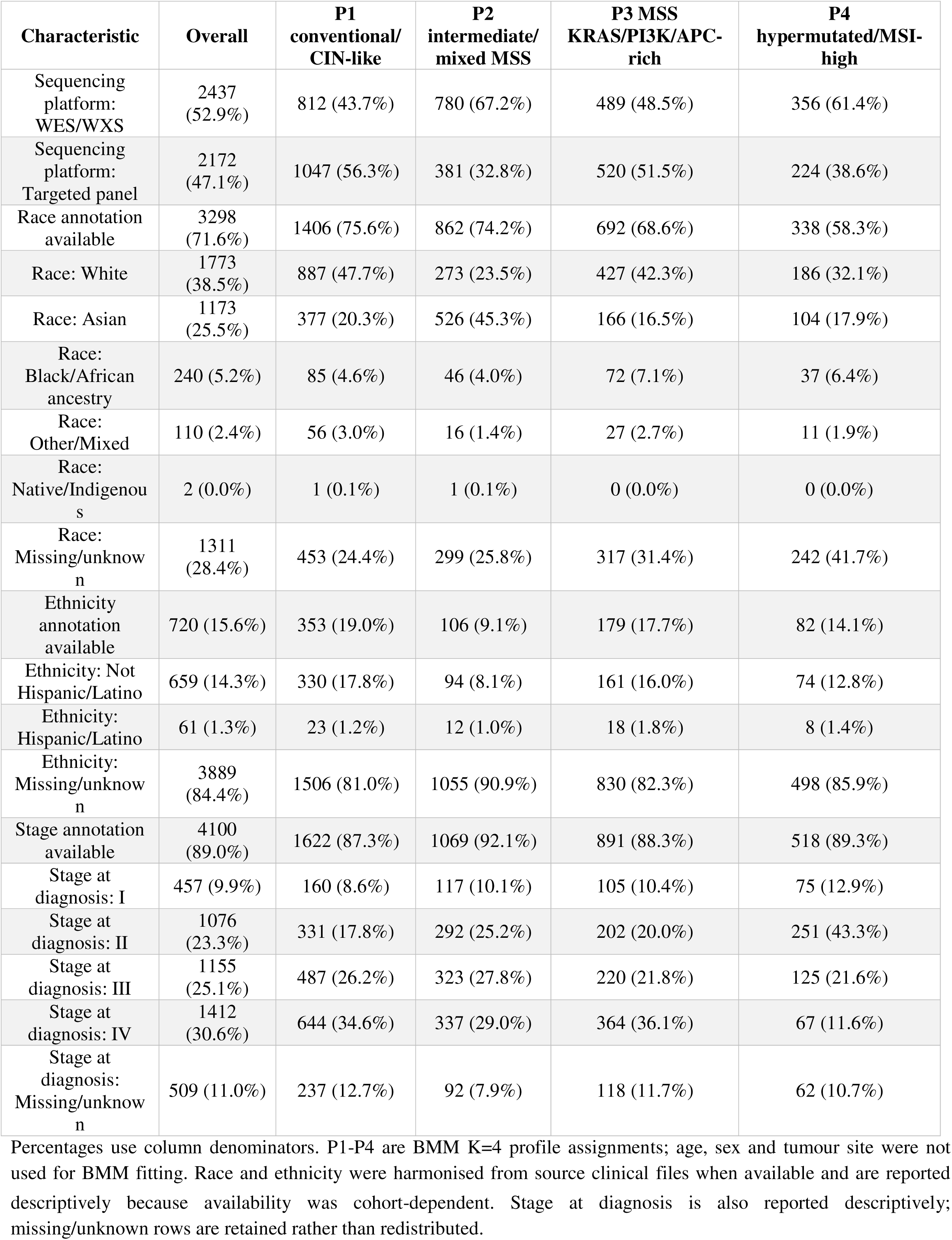
Patient characteristics of the harmonised discovery cohort by BMM K=4 profile.

**Supplementary Table 3.**
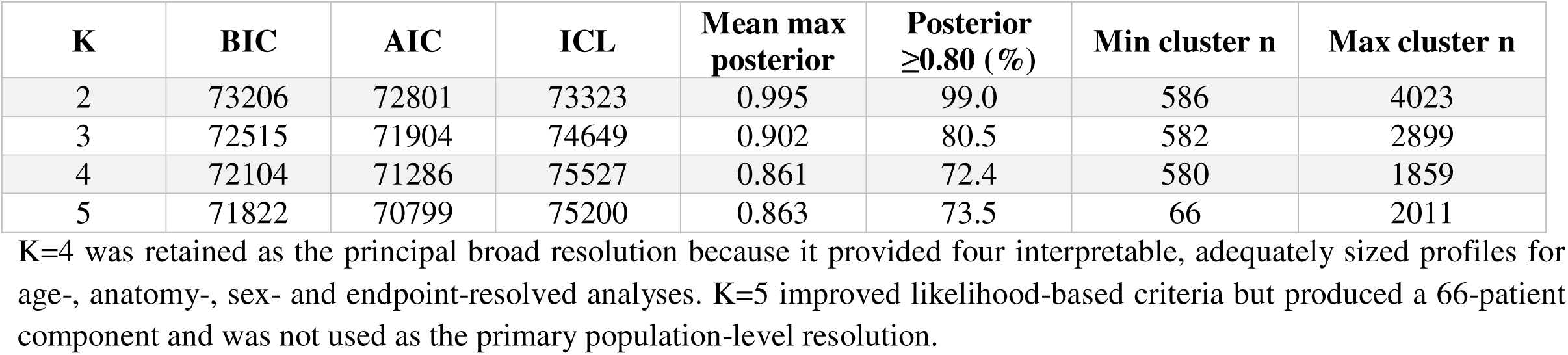
BMM model-selection and assignment diagnostics.

**Supplementary Table 4.** Full molecular-feature prevalence matrix of the BMM K=4 profiles.

| Molecular feature | P1 conventional/<br>CIN-like<br>(n=1,859) | P2 intermediate/mixed<br>MSS<br>(n=1,161) | P3 MSS<br>KRAS/PI3K/APC-rich<br>(n=1,009) | P4 hypermutated/MSI-high<br>(n=580) |
| --- | --- | --- | --- | --- |
| MSI-H/dMMR | 0.35 | 1.78 | 2.20 | 84.97 |
| FGA-high | 69.91 | 50.96 | 31.84 | 6.34 |
| TMB-high | 2.96 | 1.55 | 13.38 | 98.79 |
| APC | 100.00 | 2.33 | 84.14 | 52.59 |
| TP53 | 88.86 | 52.63 | 38.26 | 28.79 |
| KRAS | 22.65 | 13.87 | 89.69 | 31.90 |
| NRAS | 6.78 | 2.50 | 0.99 | 5.34 |
| BRAF | 3.60 | 13.70 | 2.18 | 41.03 |
| RNF43 | 0.43 | 7.84 | 1.09 | 46.90 |
| PIK3CA | 7.15 | 6.03 | 41.13 | 43.45 |
| PTEN | 0.70 | 2.33 | 8.03 | 27.41 |
| SMAD4 | 7.85 | 9.04 | 22.79 | 15.17 |
| SMAD2 | 1.08 | 0.95 | 9.71 | 11.21 |
| TGFBR2 | 0.32 | 0.95 | 2.58 | 24.14 |
| ACVR2A | 0.11 | 0.78 | 3.87 | 28.10 |
| FBXW7 | 10.27 | 3.19 | 21.31 | 36.21 |
| SOX9 | 5.86 | 3.62 | 19.72 | 20.34 |
| CTNNB1 | 1.40 | 4.91 | 6.44 | 22.41 |
| TCF7L2 | 10.01 | 1.72 | 13.78 | 28.62 |
| ARID1A | 4.25 | 3.62 | 6.24 | 48.62 |
| KMT2D | 2.90 | 3.19 | 4.56 | 58.62 |
| ATM | 3.87 | 2.93 | 7.73 | 37.07 |
| ERBB2 | 2.47 | 1.89 | 1.78 | 18.28 |

| Molecular feature | P1 conventional/<br>CIN-like<br>(n=1,859) | P2<br>intermediate/mixed<br>MSS<br>(n=1,161) | P3 MSS<br>KRAS/PI3K/APC-rich<br>(n=1,009) | P4 hypermutated/MSI-high<br>(n=580) |
| --- | --- | --- | --- | --- |
| GNAS | 0.65 | 0.69 | 4.46 | 10.52 |
| POLE | 1.24 | 0.69 | 2.78 | 36.21 |
| POLD1 | 0.65 | 1.21 | 1.59 | 25.34 |
| MLH1 | 0.32 | 1.03 | 0.30 | 19.83 |
| MSH2 | 0.54 | 0.34 | 0.10 | 19.66 |
| MSH6 | 1.08 | 0.69 | 0.50 | 26.38 |
| PMS2 | 0.05 | 0.60 | 0.79 | 12.76 |
| EPCAM | 0.05 | 0.09 | 0.20 | 3.10 |

**Notes.** Values are percentages from the frozen K=4 feature-prevalence matrix. Age, sex and tumour site were not used in BMM fitting. TMB-high was defined as TMB $\geq 10$ mutations/Mb. FGA-high was defined within each study as FGA ≥ the study-specific median among FGA-evaluable patients after age filtering. MSI-H/dMMR was derived from harmonised study-provided MSI/MMR fields. Gene-level indicators represent qualifying functional somatic alterations.
**MSS/pMMR.** MSS/pMMR is the complement of MSI-H/dMMR among MSI/MMR-evaluable tumours and was not an additional independent BMM input. Derived MSS/pMMR prevalences were 99.65% (P1), 98.22% (P2), 97.80% (P3) and 15.03% (P4).
**Abbreviations.** BMM, Bernoulli mixture model; CIN, chromosomal instability; FGA, fraction genome altered; MSI-H/dMMR, microsatellite instability-high/deficient mismatch repair; MSS/pMMR, microsatellite stable/proficient mismatch repair; TMB, tumour mutational burden.

**Supplementary Table 5.** Full linear and quadratic age-trend statistics for Figure 1E profiles.

| Profile | Selected displayed term | Linear p | Linear q (8-test BH) | Quadratic p | Quadratic q (8-test BH) | Study+sex adjusted linear p | Study+sex adjusted quadratic p | Peak decade | Trough decade |
| --- | --- | --- | --- | --- | --- | --- | --- | --- | --- |
| P1 | quadratic | 9.4e-10 | 3.7e-09 | 4.2e-06 | 8.5e-06 | 0.001 | 8.3e-06 | 40-49 | <30 |
| P2 | linear | 0.067 | 0.09 | 0.836 | 0.836 | 0.011 | 0.099 | <30 | 80+ |
| P3 | linear | 3.2e-05 | 5.1e-05 | 0.245 | 0.28 | 5.6e-05 | 0.076 | 80+ | <30 |
| P4 | quadratic | 1.5e-09 | 3.9e-09 | 8.4e-15 | 6.7e-14 | 0.018 | 2.3e-09 | 80+ | 50-59 |

*The main figure displays the biologically selected term for each profile, but both linear and quadratic terms are reported here to avoid post-hoc ambiguity. q values are BH-adjusted across the eight unadjusted profile trend tests.*

**Supplementary Table 6.** Patient-level study-, sex- and site-adjusted trend sensitivity models.

| Profile | Selected term | Study+sex+site adjusted linear p | Study+sex+site adjusted quadratic p | Selected adjusted q (4-test BH) |
| --- | --- | --- | --- | --- |
| P1 | quadratic | 0.936 | 4.0e-04 | 5.4e-04 |
| P2 | linear | 0.001 | 0.084 | 0.001 |
| P3 | linear | 1.2e-04 | 0.031 | 2.3e-04 |
| P4 | quadratic | 0.515 | 6.9e-07 | 2.8e-06 |

**Supplementary Table 7.** Study-level random-effects heterogeneity summary. *Two-stage analyses were used as sensitivity analyses to quantify study-level heterogeneity. Anatomical effects are random-effects odds ratios for right colon versus left/distal colon. Age-trend effects are random-effects coefficients for the selected* Figure 1E *term. I^2^ quantifies the percentage of observed effect variability attributable to between-study heterogeneity*.

| Analysis | Profile | Effect tested | Random-effects estimate (95% CI) | P value | $I^2$ | Interpretation |
| --- | --- | --- | --- | --- | --- | --- |
| Anatomical enrichment | P1 | right vs left OR | OR 0.26 (0.20-0.36) | 3.9e-18 | 50% | left/distal-enriched |
| Anatomical enrichment | P2 | right vs left OR | OR 0.87 (0.72-1.06) | 0.176 | 10% | intermediate/not clearly side-enriched |
| Anatomical enrichment | P3 | right vs left OR | OR 2.01 (1.65-2.44) | 2.5e-12 | 5% | right-shifted |
| Anatomical enrichment | P4 | right vs left OR | OR 6.03 (4.67-7.80) | 5.5e-43 | 0% | strongly right-enriched |
| Age-trend coefficient | P1 | quadratic | beta -0.10 (-0.13 to -0.07) | 2.9e-09 | 0% | robust quadratic pattern |
| Age-trend coefficient | P2 | linear | beta -0.11 (-0.18 to -0.03) | 0.005 | 25% | modest cohort-sensitive linear decrease |
| Age-trend coefficient | P3 | linear | beta 0.14 (0.07 to 0.20) | 4.8e-05 | 11% | robust linear increase |
| Age-trend coefficient | P4 | quadratic | beta 0.07 (-0.01 to 0.16) | 0.082 | 66% | heterogeneous age-shape |

## Supplementary figure legends

Supplementary Figure 1. Bernoulli mixture model validation and cohort composition. A. Information criteria across K. B. Assignment confidence across K. C. K=4 profile sizes. D. Profile composition by study. AIC, BIC and the reported positive-form ICL values are shown as minimisation criteria. K=4 was retained as the principal broad resolution while cohort composition and assignment confidence are shown explicitly.

Supplementary Figure 2. BMM-CMS molecular-profile proximity independent of anatomical side. Pearson correlations between P1-P4 and CMS1-CMS4 across eight shared molecular features, with nominal p and BH-FDR q values. Anatomical side was omitted from the feature vectors to evaluate molecular similarity independently of tumour location (left). Euclidean distance after feature-wise z-scoring; lower values indicate greater proximity (right). P1 aligns with CMS2, P3 with CMS3 and P4 with CMS1, whereas P2 shows weak/partial proximity to CMS4. This represents a general feature-vector comparison, not patient-level CMS classification.

Supplementary Figure 3. Age-trend robustness after study omission and covariate adjustment. A. Leave-one-study-out significance retention for the selected Figure 1E trend terms. B. Study-, sex- and site-adjusted q values for the same terms. P1, P3 and P4 show the strongest leave-one-study-out retention; P2 is significant after adjustment but less consistent across study omissions.

Supplementary Figure 4. Study-level heterogeneity of selected Figure 1E age-trend coefficients. Forest plots show study-specific and random-effects pooled coefficients for P1 quadratic, P2 linear, P3 linear and P4 quadratic terms. P1, P2 and P3 are supported by two-stage random-effects analysis, whereas the P4 age-shape magnitude is more heterogeneous across studies.

Supplementary Figure 5. Study-level heterogeneity of anatomical profile enrichment. Forest plots show study-specific and random-effects pooled estimates for right/proximal versus left/distal enrichment of each profile. P1 is left/distal-enriched, P3 is right-shifted and P4 strongly right-enriched; P2 remains intermediate.

Supplementary Figure 6. Platform sensitivity of BMM K=4 age trajectories and molecular profile architecture. A. Age-decade prevalence of fixed principal K=4 assignments in WES/WXS and targeted-panel cohorts, with age-by-platform interaction tests. B. Within-profile prevalence of the 31 BMM inputs across platforms, with concordance and mean absolute difference. FGA-high is highlighted because evaluability differed markedly. Assignments were not refitted.

Supplementary Figure 7. BMM K=4 sensitivity refit after excluding FGA-high. A. Original and 30-feature refitted age-decade trajectories. B. Molecular-feature prevalence within original and refitted profiles across retained inputs. Overall assignment retention was 96.4% (adjusted Rand index 0.903; normalised mutual information 0.882). Information criteria were not compared across models fitted to different feature dimensions.

Supplementary Figure 8. MLTA and GAMM analyses reproduce the molecular profiles and age-prevalence structure identified by BMM. A. Concordance between BMM and MLTA molecular centroids. Each point represents one molecular feature within one matched profile (31 features × four profiles); colours denote P1-P4 and the dashed line indicates identity. The overall Pearson correlation was 0.995, with a mean absolute error of 1.1 percentage points. B. Row-normalised cross-classification of patient-level assignments. Cells show the percentage and number of patients within each BMM profile assigned to each mapped MLTA profile; 91.2% of assignments matched and the adjusted Rand index was 0.78. C. Percentage of decade-resolved profile-composition deviance retained by the conventional <50/≥50 comparison. Points show observed estimates and bars show study-stratified bootstrap 95% confidence intervals from 2,000 resamples. The binary split retained 17.8% (95% confidence interval, 7.3-29.9%) of decade-resolved deviance under BMM and 21.4% (10.5-33.3%) under MLTA. D. Prevalence of each matched molecular profile across age decades under BMM (solid lines and circles) and MLTA (dashed lines and squares). Decade-resolved prevalence trajectories were strongly concordant overall (Pearson r=0.953) and within each profile (r=0.979-0.999). E. GAMM analysis of continuous age. For each profile, a binomial GAMM containing a cubic-spline age function and a study-level random intercept was compared with a simpler linear-age mixed model. Curves and shaded areas show model estimates and approximate 95% confidence intervals over ages 25-85 years (4,528 patients); open symbols show observed decade prevalence. P1 and P4 display GAMM estimates because non-linear spline components were supported under both BMM and MLTA. P2 and P3 display the parsimonious linear mixed-model estimates because both had significant linear trends (P2 q=0.011 under both methods; P3 q=5.59×10 under BMM and 3.41×10 under MLTA), whereas adding non-linear spline terms did not improve either trajectory (P2 q=0.342 and 0.332; P3 q=0.802 under both methods). Age was available for 4,607 of 4,609 patients.

Supplementary Figure 9. Age-decade prevalence of tumour mutational burden, copy-number alteration and mismatch-repair features in the pooled CRC cohort. The four panels show directly observed patient-level prevalence across age decades after combining all BMM profiles. A. TMB-high, defined as tumour mutational burden ≥10 mutations/Mb. B. FGA-high, defined relative to the within-study median fraction genome altered among FGA-evaluable patients; C. MSI-H/dMMR; and D. MSS/pMMR/MSI-L, defined as the complementary category among tumours with evaluable MSI/MMR status. Points denote the observed prevalence within each age decade and error bars show Wilson 95% confidence intervals. The marker-specific evaluable sample size is shown for each decade; denominators therefore vary according to data availability. Quadratic age effects were tested using nested likelihood-ratio tests comparing linear-age models with linear-plus-quadratic-age models. Reported p values test the addition of the quadratic age term, and q values were obtained by Benjamini-Hochberg correction across the four feature-specific tests. TMB-high and MSI/MMR status showed strong U-shaped age patterns, whereas the directly measured FGA-high trajectory did not meet the corrected significance threshold. Total evaluable populations were 4,607 patients for TMB-high, 2,379 for FGA-high and 4,160 for MSI-H/dMMR and MSS/pMMR/MSI-L.

Supplementary Figure 10. P1 retains an APC/MSS backbone across age extremes but shifts in TP53 and anatomy. Selected features are compared within P1 between <30 and 80+ years and between <40 and ≥70 years. Points show older-minus-younger prevalence differences with Newcombe 95% confidence intervals; q values are BH-FDR-adjusted Fisher exact tests. FGA-high uses available-case denominators.

Supplementary Figure 11. P4 age extremes share a hypermutated state but differ in accompanying molecular and anatomical contexts. Selected features are compared within P4 between <30 and 80+ years and between <40 and ≥70 years. Older P4 is BRAF/RNF43/right-sided enriched; younger P4 is APC/KRAS/somatic-MMR-alteration enriched. POLE/POLD1 is not significantly enriched at either extreme. These proxies do not establish germline status, MLH1 methylation or CIMP.

Supplementary Figure 12. Molecular profiles remain recognisable across age, but decade-resolved redistribution is poorly captured by age 50. A. Within each profile, prevalence of all 31 BMM inputs in each decade versus profile-wide prevalence. B. The same inputs compared between <50 and ≥50 patients. C. Absolute age-50 prevalence difference versus full decade peak-to-trough range. D. Likelihood-ratio deviance partitioned into the component represented by age 50 and additional decade-resolved structure within the pooled groups.

Supplementary Figure 13. Raw overall survival by BMM K=4 profile. Kaplan-Meier curves show unadjusted overall survival for P1-P4 with numbers at risk. The global log-rank test compares profiles. This analysis is a descriptive clinical benchmark rather than an adjusted prognostic model.

Supplementary Figure 14. Rectal cancer profiles. A. Profile composition across left/distal colon, rectum and right/proximal colon. B. Rectum-minus-comparator profile-prevalence differences. C. Rectum-only age-decade trajectories with confidence ribbons.

Supplementary Figure 15. Rectal profile composition after excluding MSK Rectal 2022. Profile composition across left/distal colon, rectum and right/proximal colon is shown before and after exclusion. The P1-high, P2-low and P3-enriched rectal state persists.

Supplementary Figure 16. Non-overlapping MSK-IMPACT validation reproduces molecular-profile identity and non-binary age redistribution. A. Validation-cohort derivation. Among 4,993 patients with a colorectal tumour record, 3,575 had a primary CRC specimen. After excluding 1,340 patients whose MSK patient identifiers matched any discovery MSK cohort and retaining one primary specimen per patient, 2,235 non-overlapping patients remained; 2,134 reported age at diagnosis and 502 were younger than 50 years. B. Comparison of discovery and validation prevalence for each of the 30 shared molecular features within each profile (120 profile-feature points). The line and shading show the fitted linear regression and its 95% confidence interval; Pearson correlations are shown overall and by profile. ACVR2A was not assayed by MSK-IMPACT and was marginalised during profile projection. C. Profile prevalence by age decade in discovery and validation. Discovery trajectories are shown as solid profile-coloured lines with 95% Wilson confidence-interval ribbons; MSK validation trajectories are shown as dashed grey lines. Correlations compare the seven-point discovery and validation trajectories for each profile; the within-profile standardised correlation across all 28 profile-decade points was r=0.904 (P=4.38×10^−11^). D. Prespecified continuous-age terms in discovery and validation: quadratic terms for P1/P4 and linear terms for P2/P3. Models were adjusted for sex, tumour site and sequencing platform group in discovery and for sex, tumour site and IMPACT panel in validation. Points and bars show coefficients and 95% confidence intervals. Displayed q values correspond to validation models and were Benjamini-Hochberg adjusted across the four prespecified tests; P2 retained a negative linear coefficient but with weaker adjusted support. E. Percentage of decade-resolved profile-composition likelihood-ratio deviance captured by every possible integer age cut-off within the observed age range. Solid black and dashed blue curves represent the discovery and MSK validation cohorts, respectively. The dotted vertical line and open circles mark age 50; filled circles mark the best-performing threshold in each cohort. Age 50 captured 17.8% of decade-resolved deviance in discovery and 6.2% in validation, whereas the best-performing thresholds occurred at age 70 in discovery (51.3%) and age 75 in validation (49.4%). Age, sex and tumour site were excluded from all profile assignments.

