## Supplementary figures and images for "Prevalence of Colorectal Cancer Molecular Profiles Is Not Captured by a Single Age Threshold"

### Supp Fig1

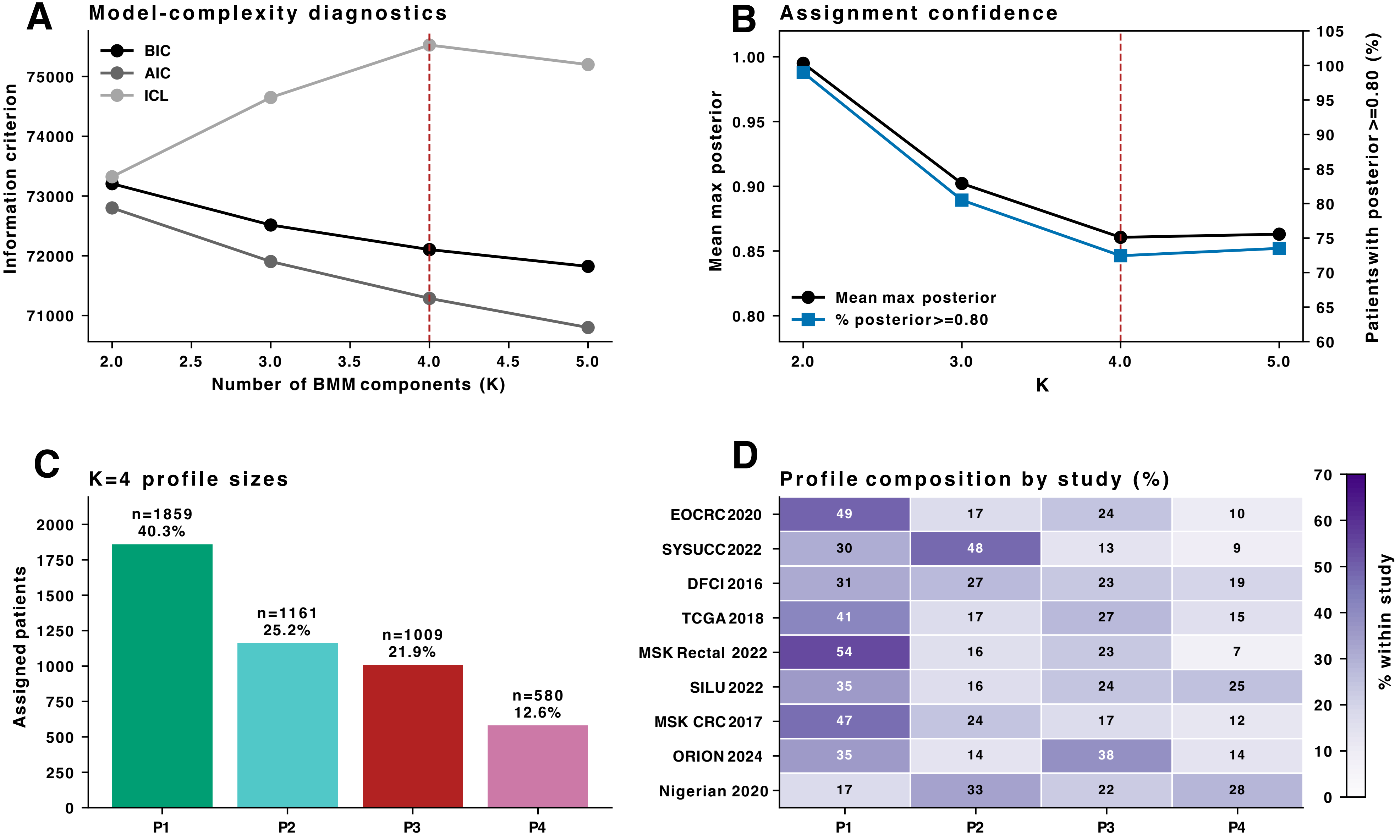

### Supp Fig2

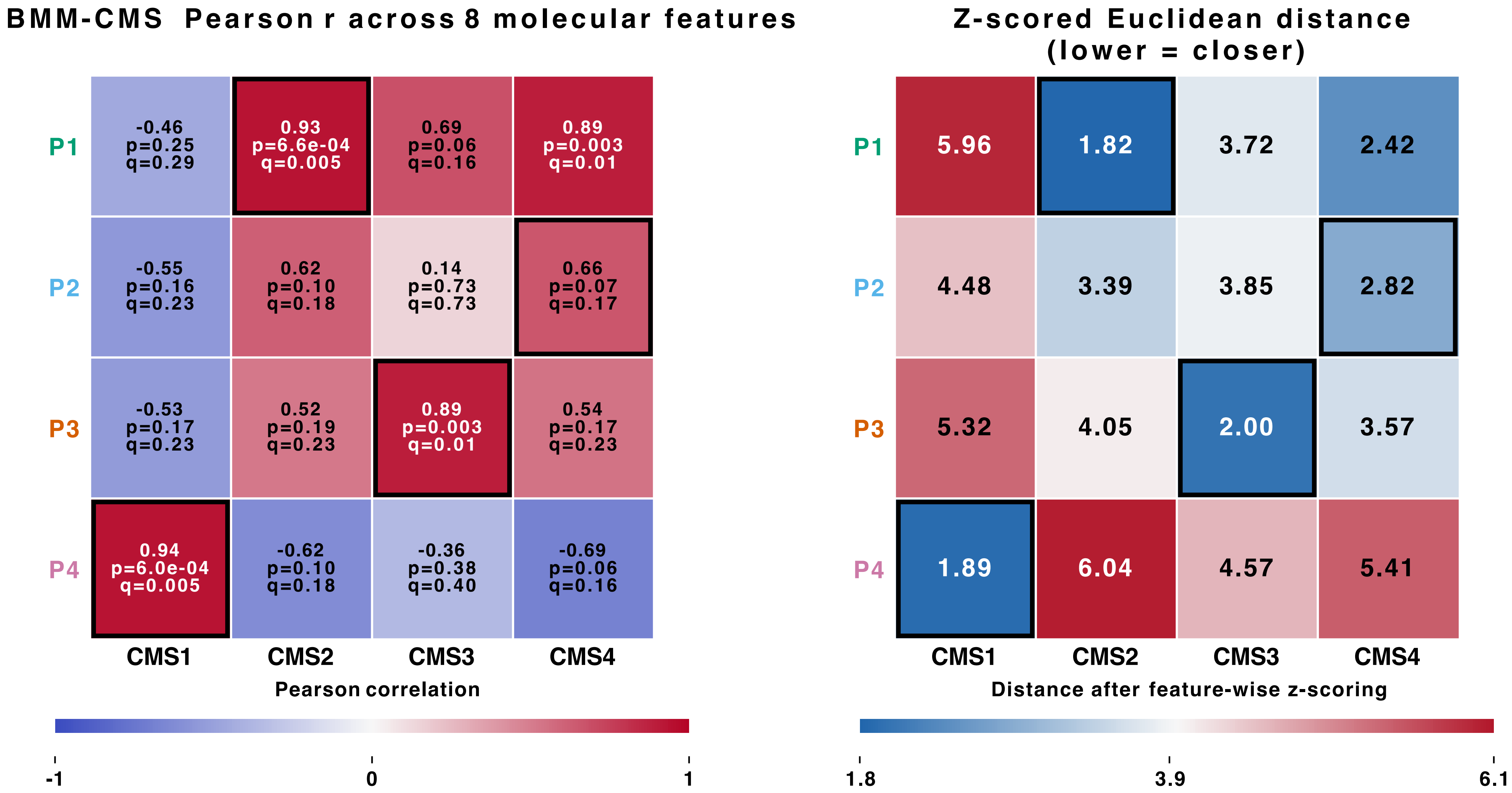

### Supp Fig3

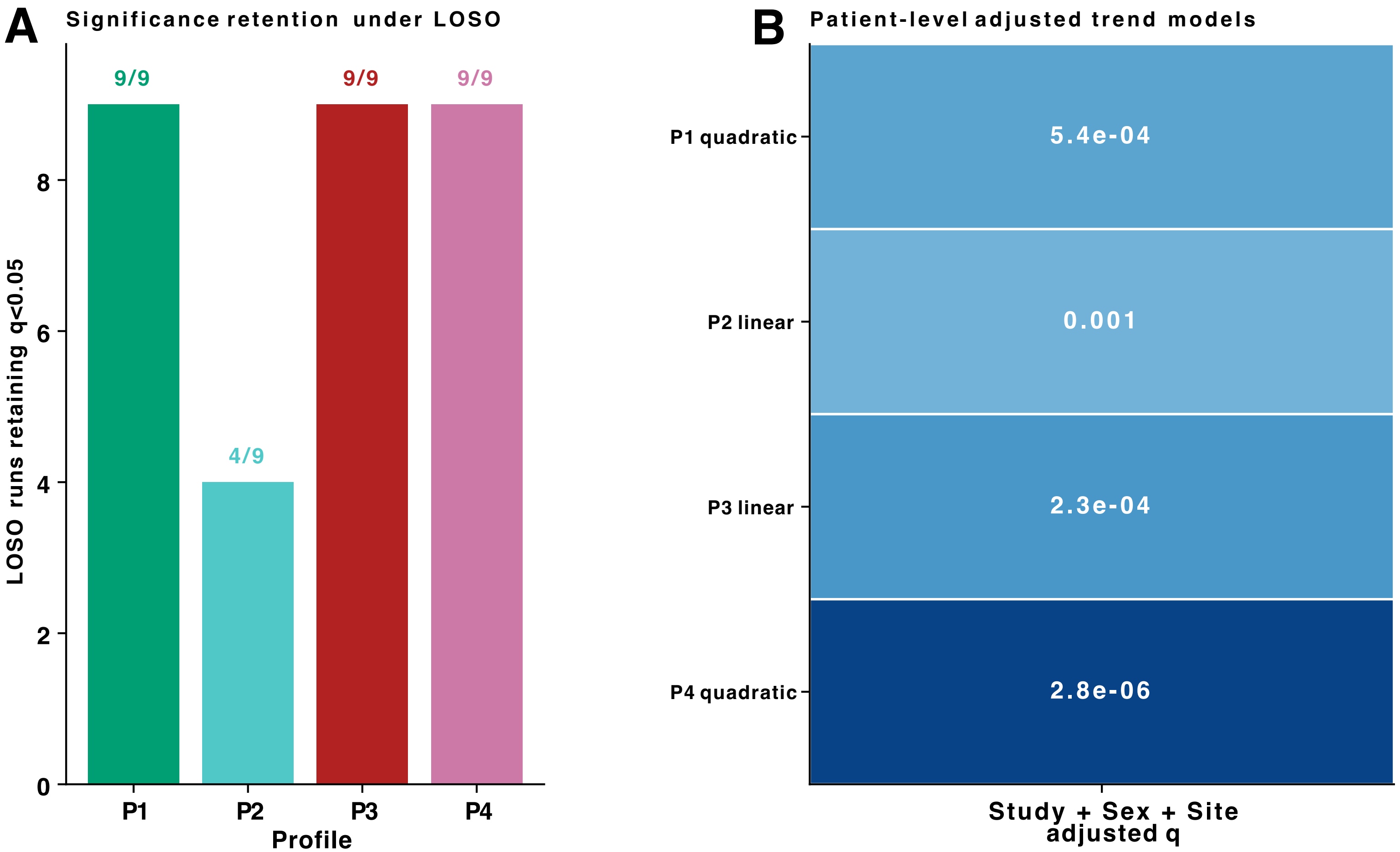

### Supp Fig4

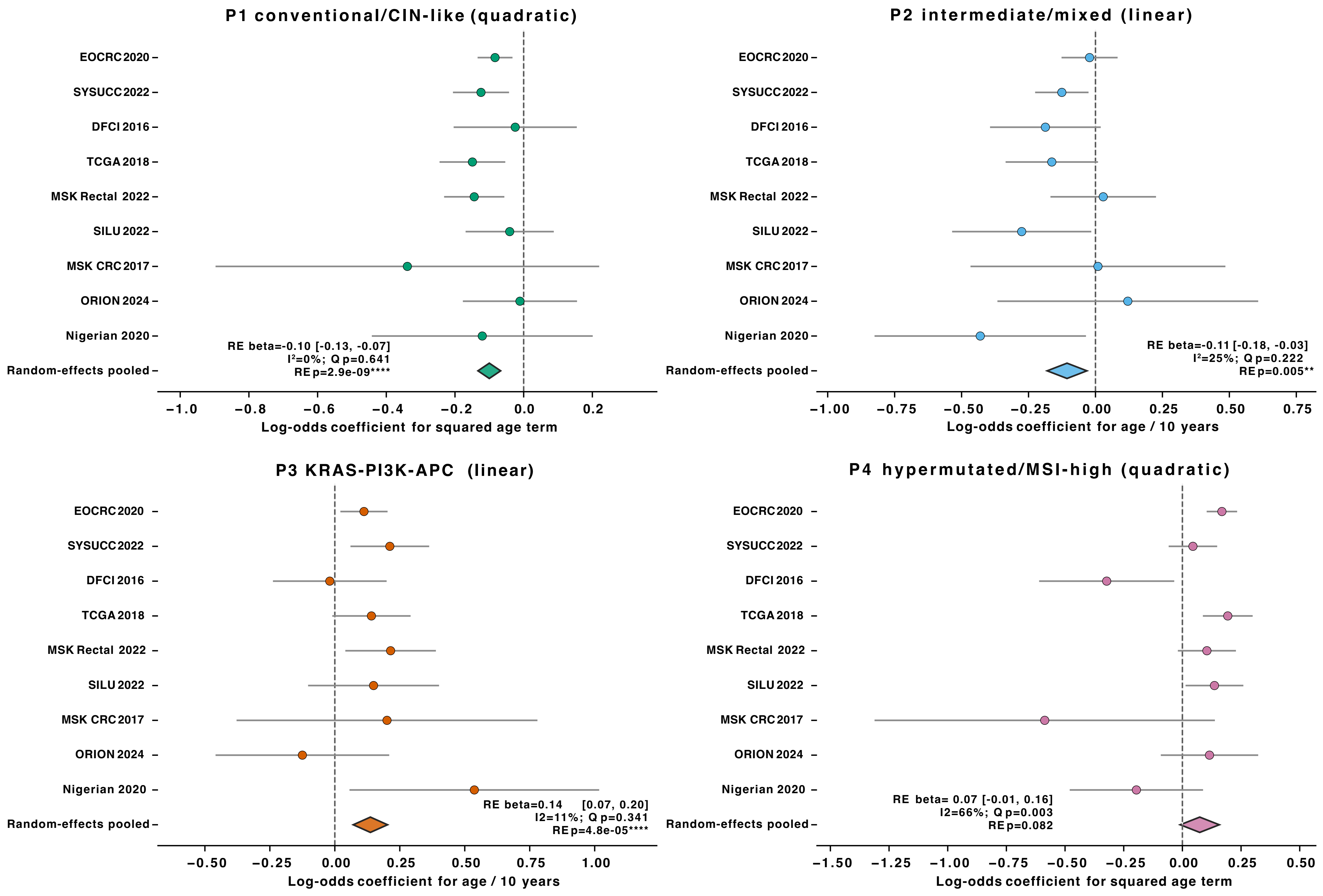

### Supp Fig5

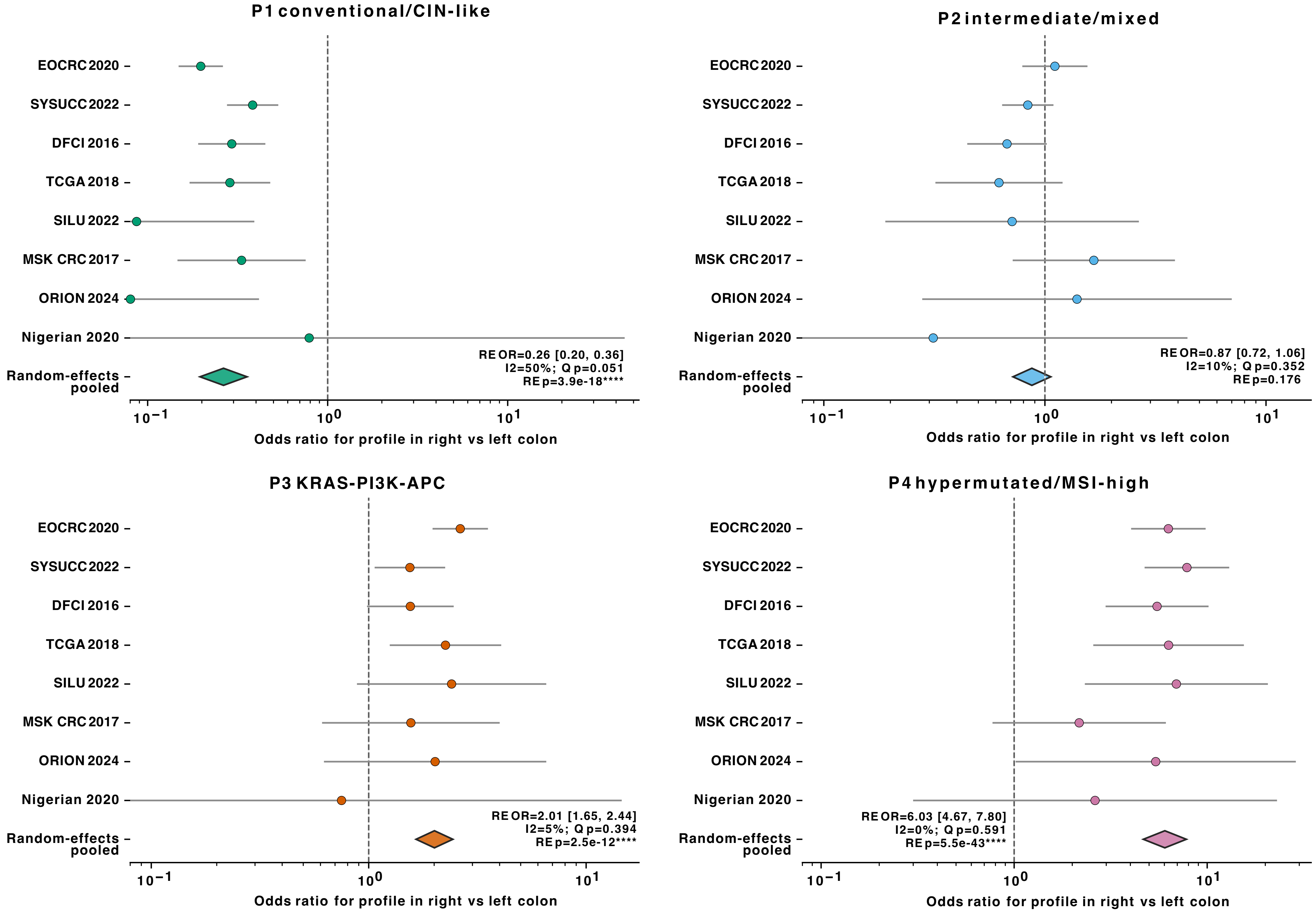

### Supp Fig6

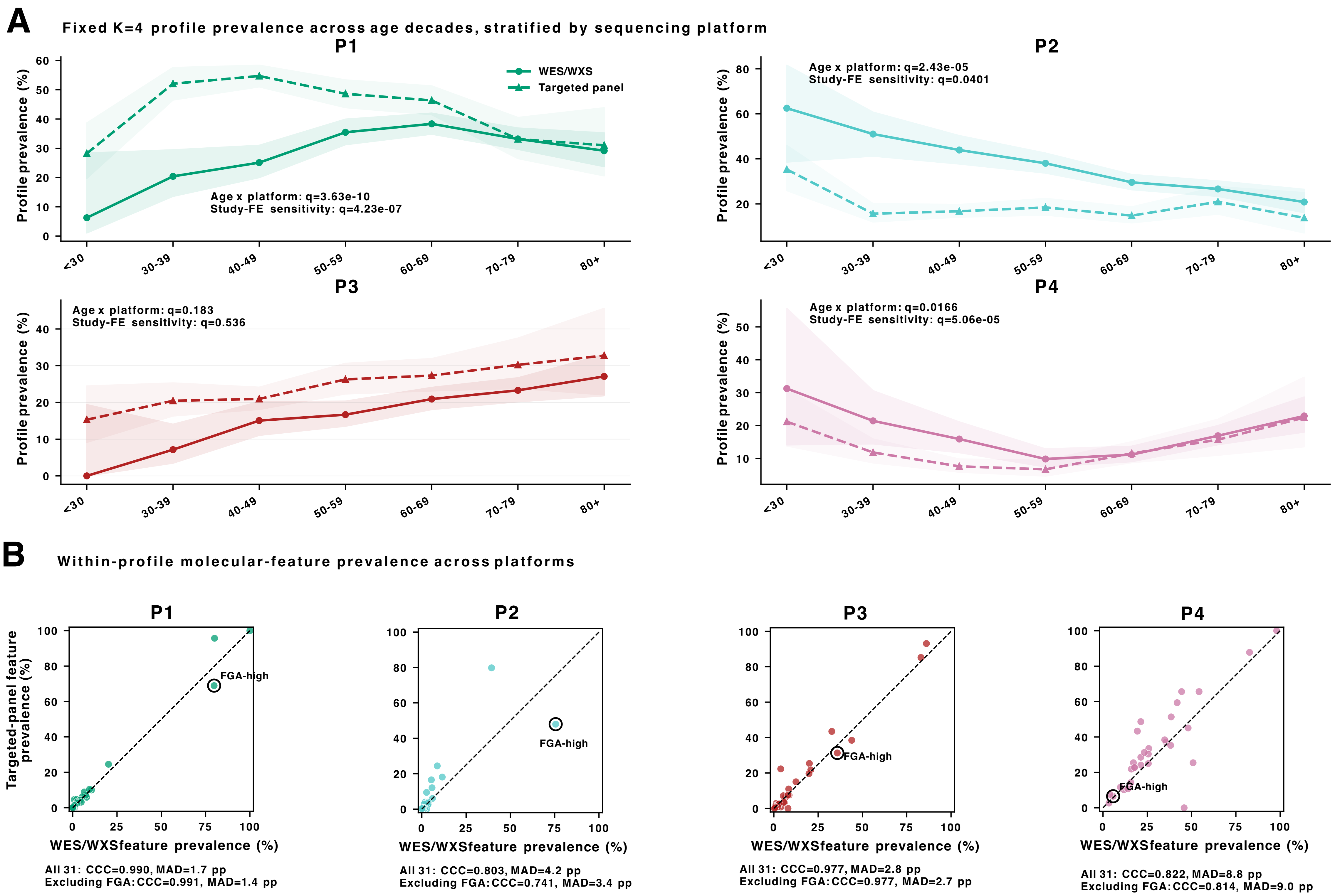

### Supp Fig7

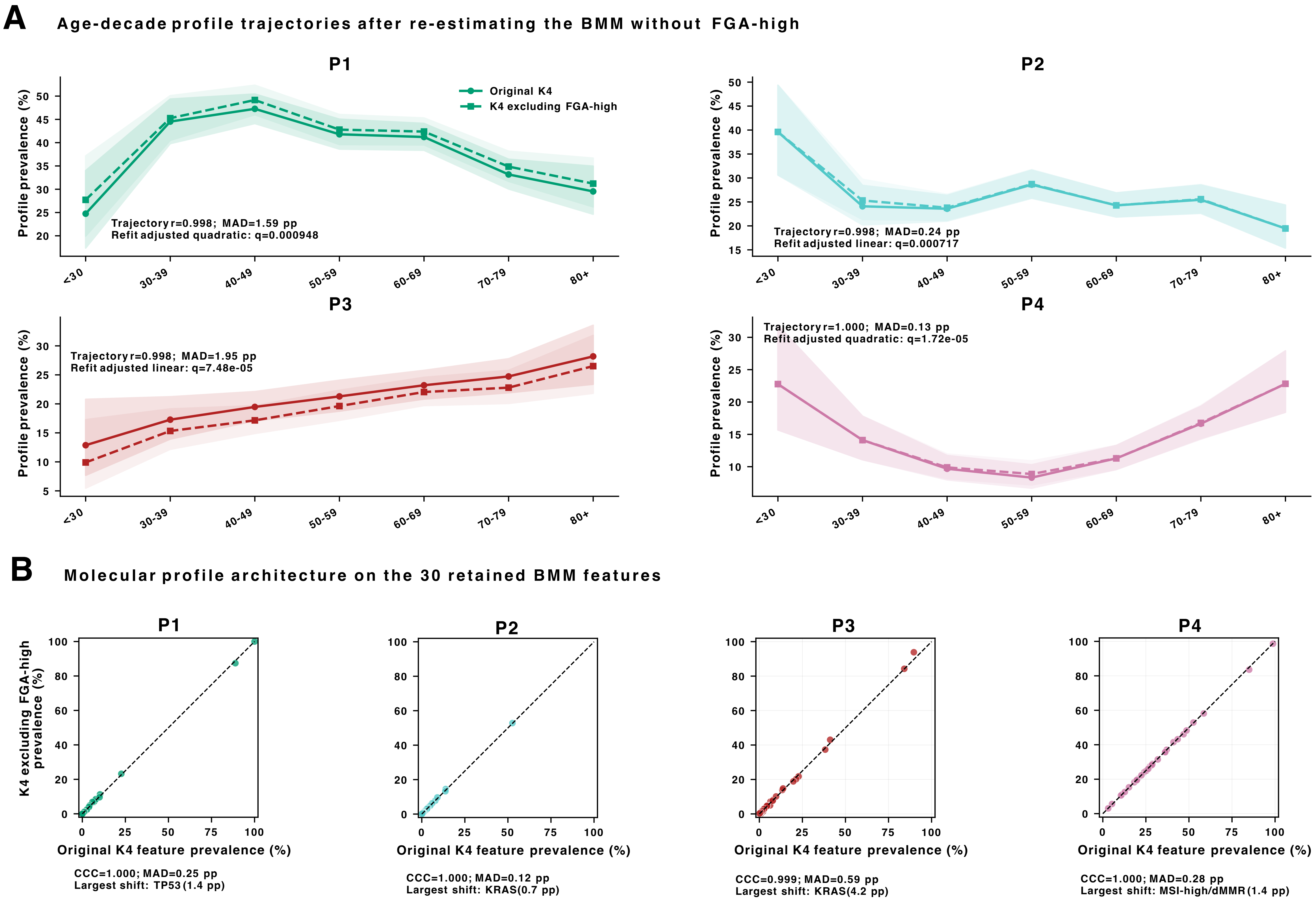

### Supp Fig8

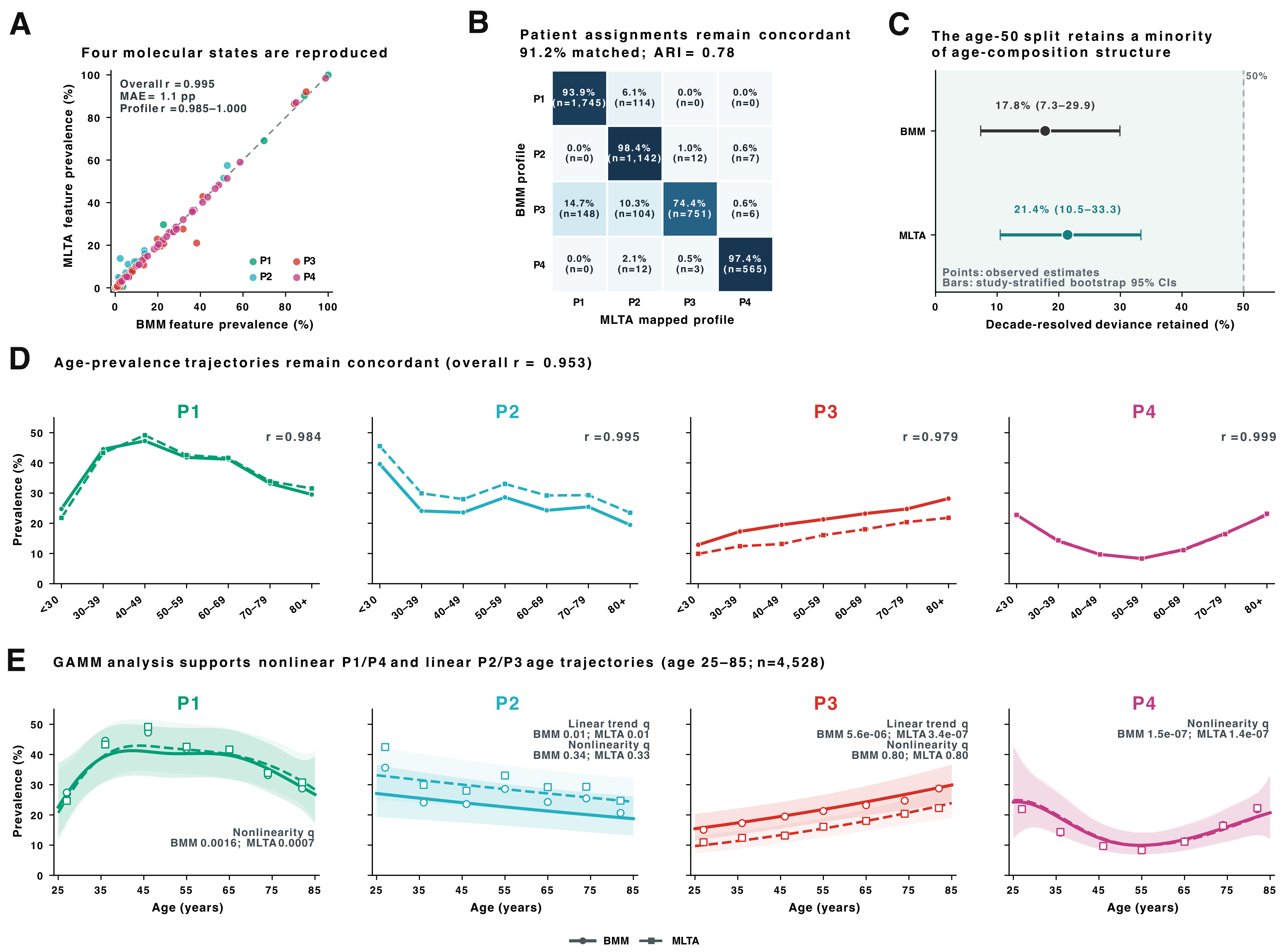

### Supp Fig9

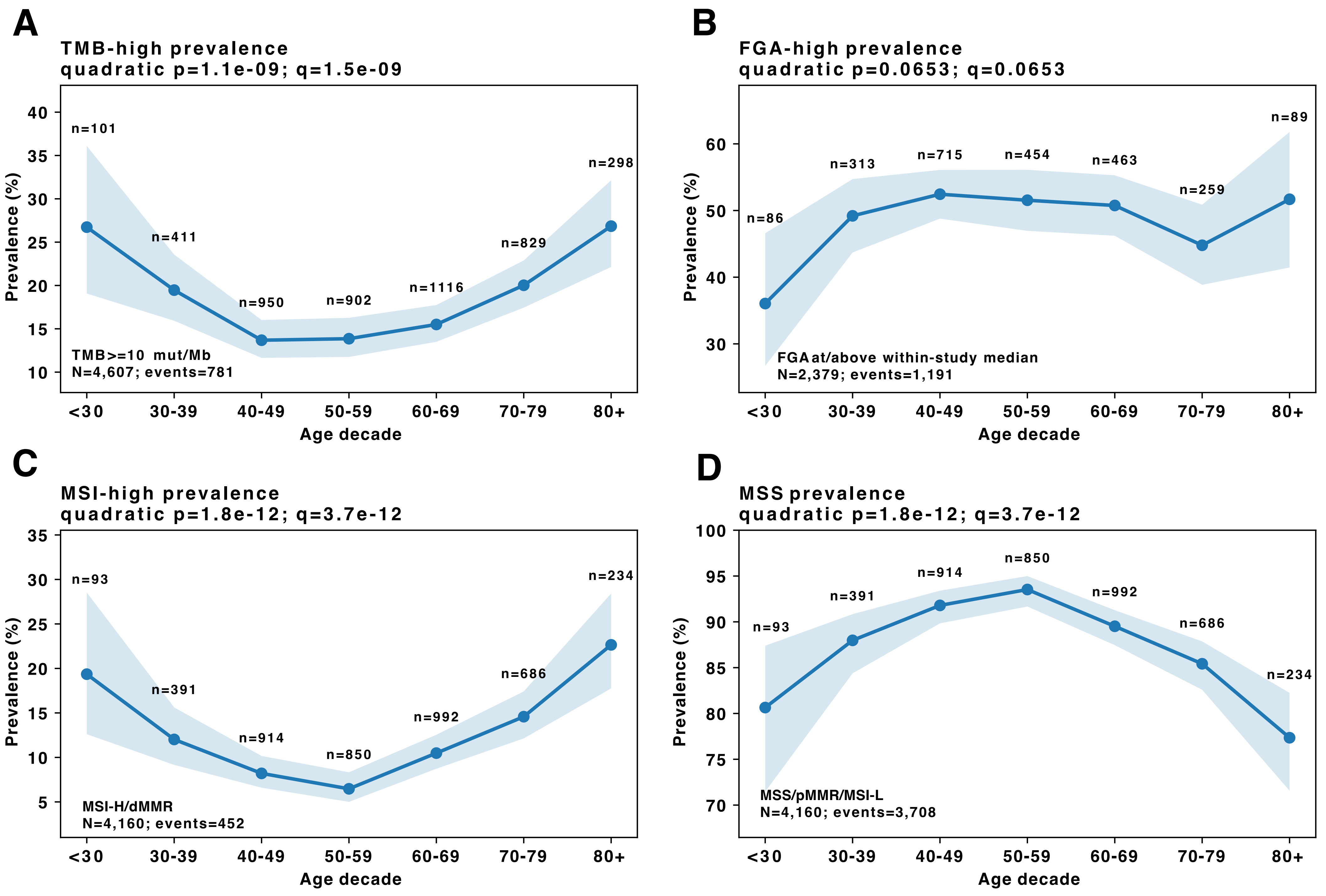

### Supp Fig10

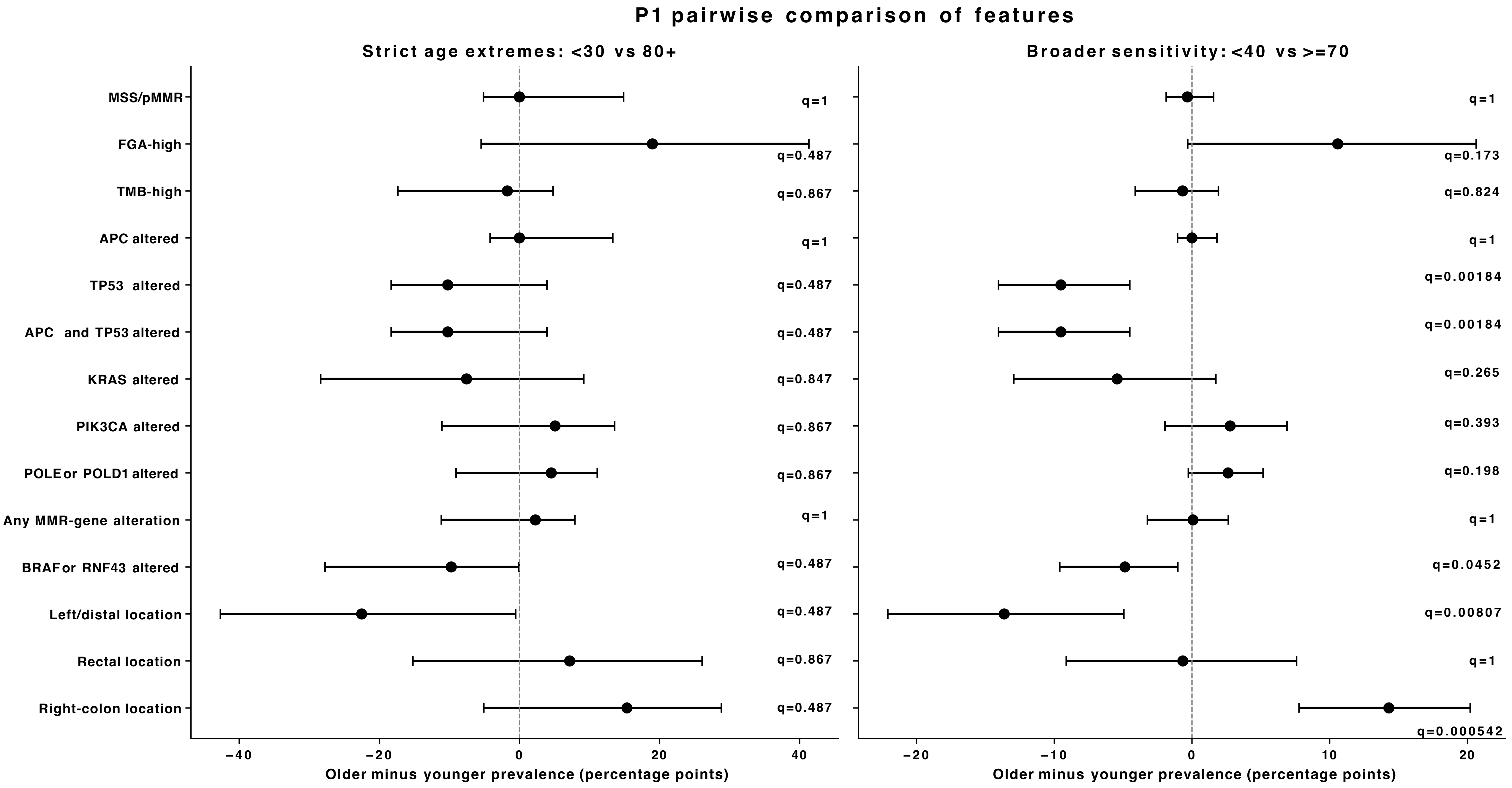

### Supp Fig11

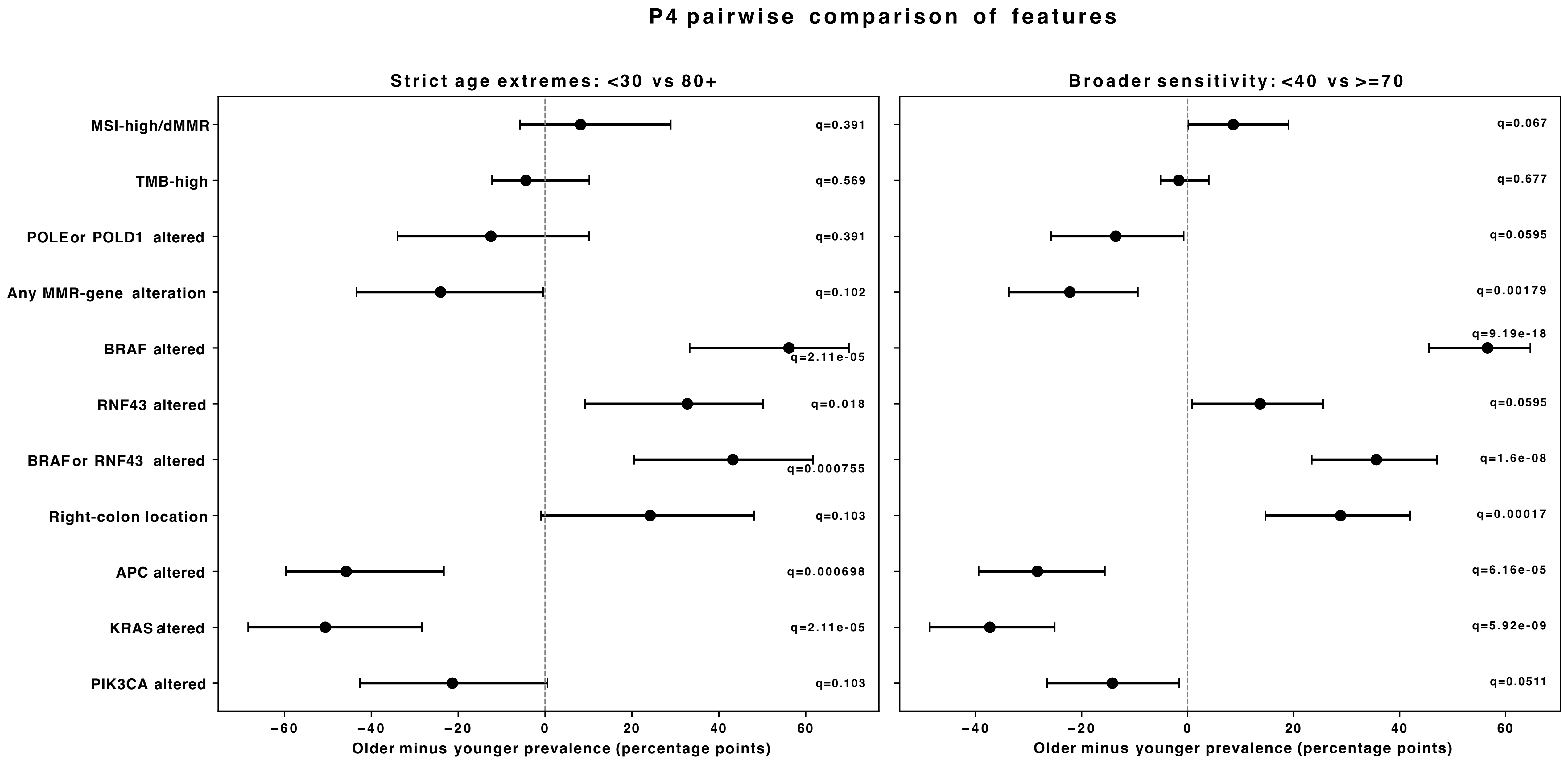

### Supp Fig12

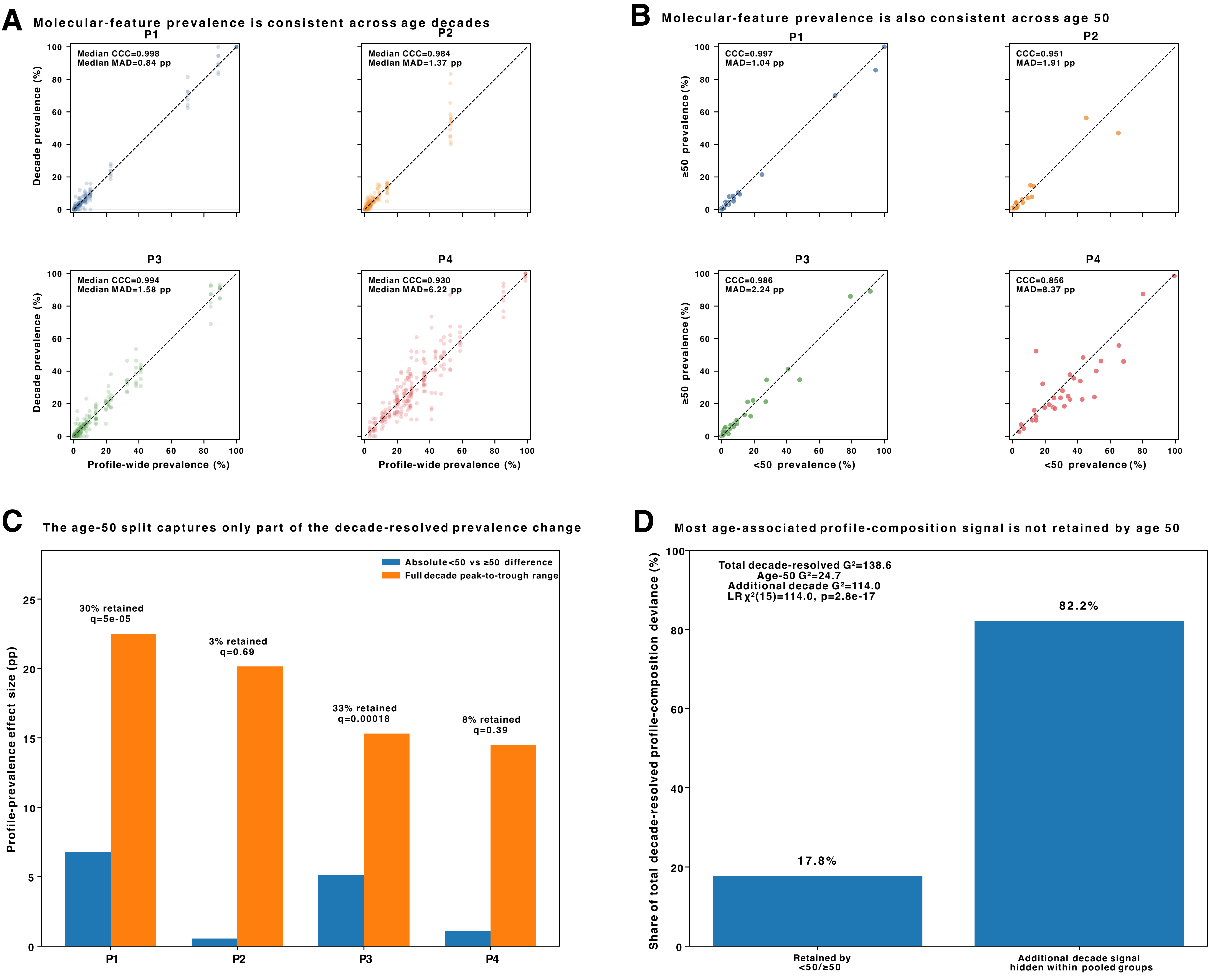

### Supp Fig13

#### Observed survival by BMM K=4 molecular profile

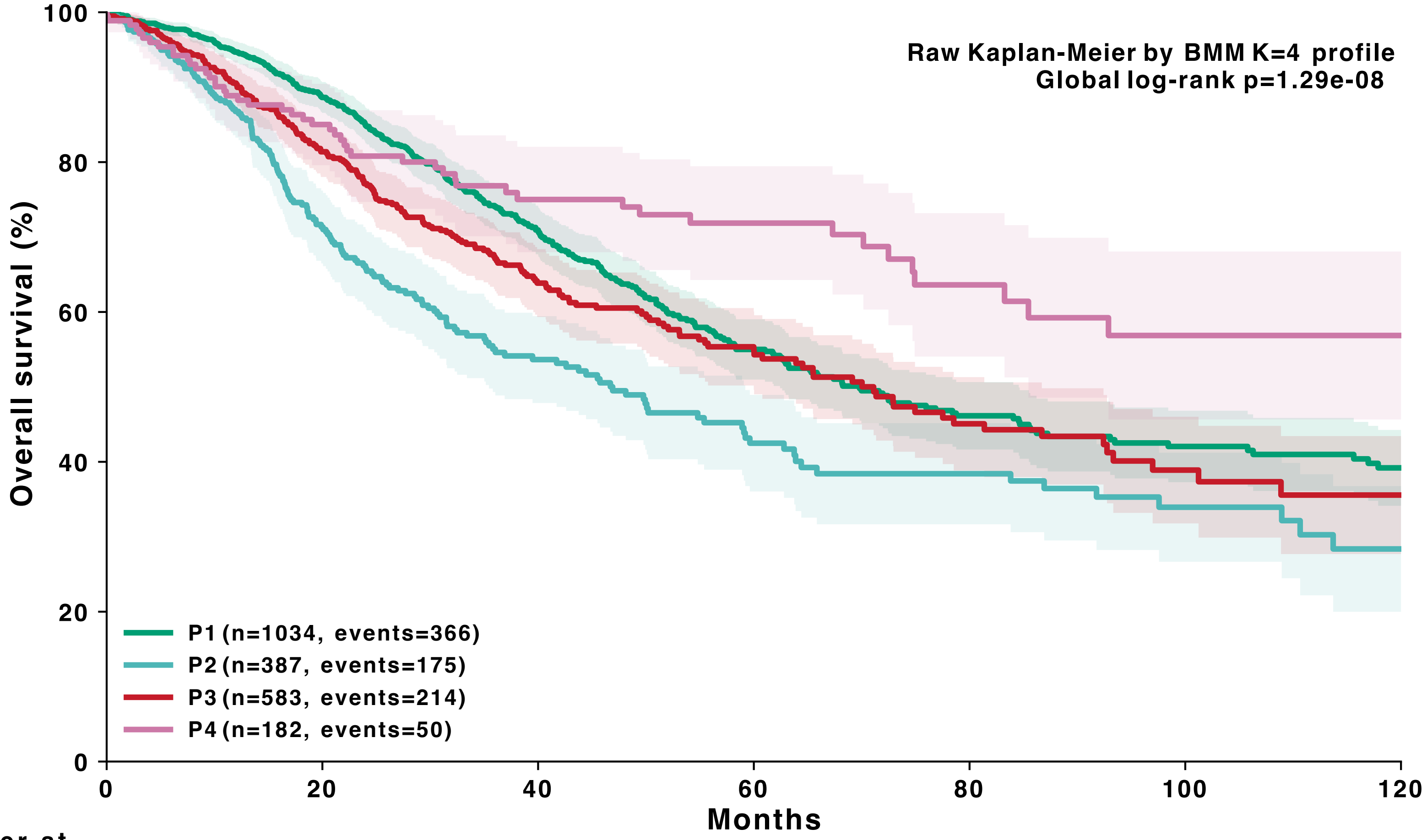


| Number at risk | 0 | 12 | 24 | 36 | 60 | 84 | 120 |
|---|---|---|---|---|---|---|---|
| P1 | 1034 | 884 | 665 | 472 | 211 | 120 | 62 |
| P2 | 387 | 295 | 183 | 124 | 61 | 39 | 13 |
| P3 | 583 | 472 | 338 | 238 | 103 | 55 | 18 |
| P4 | 182 | 146 | 111 | 86 | 53 | 28 | 15 |

### Supp Fig14

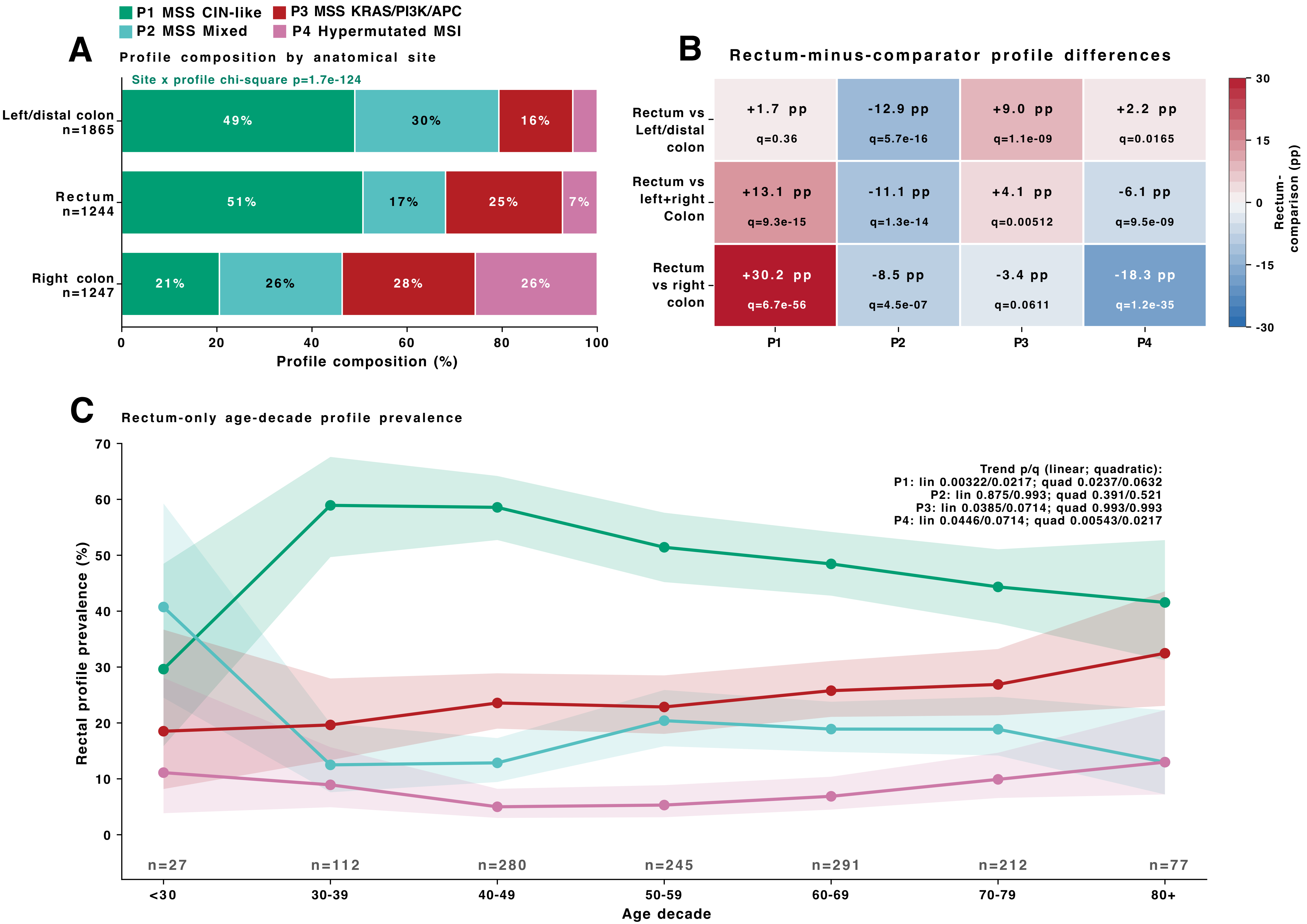

### Supp Fig15

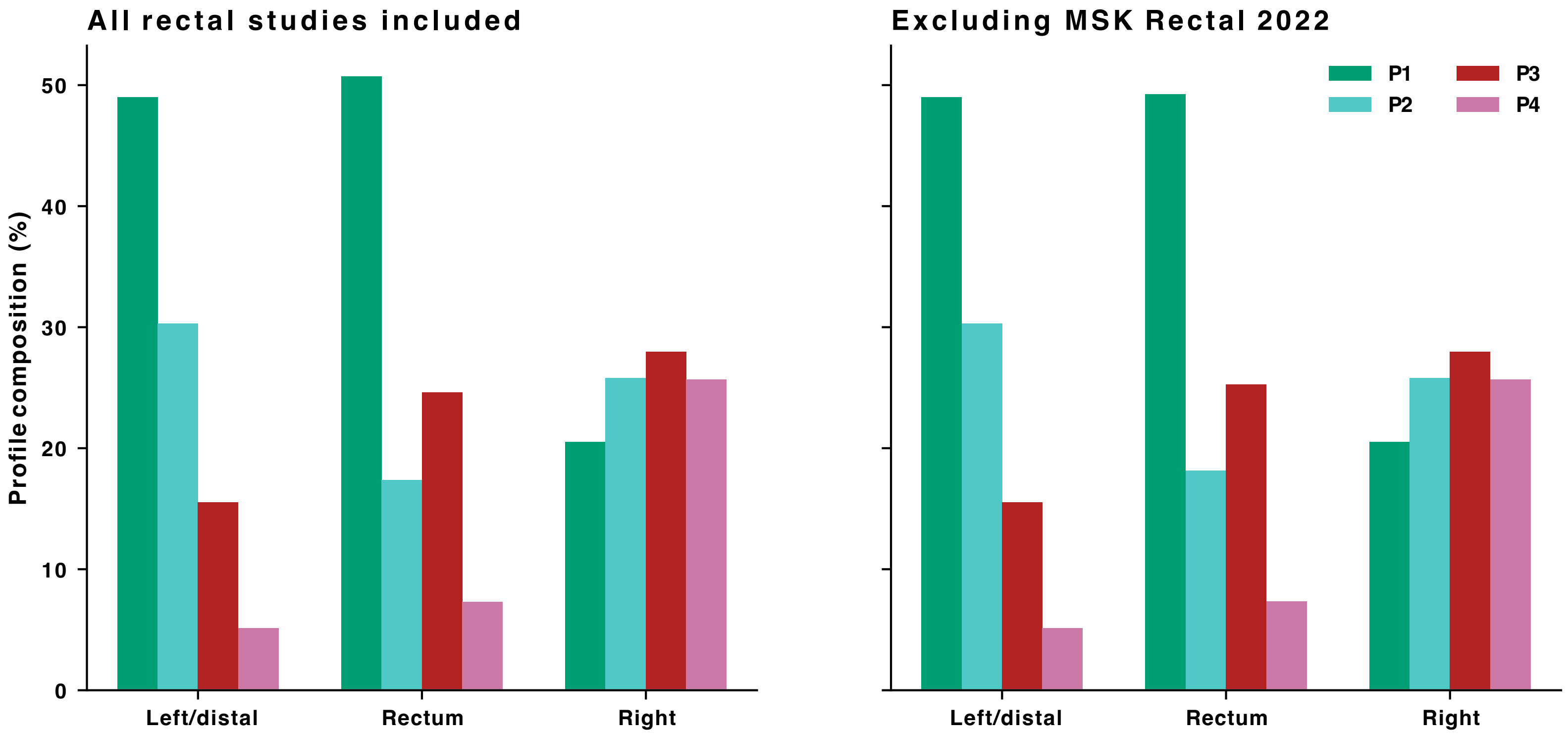

### Supp Fig16

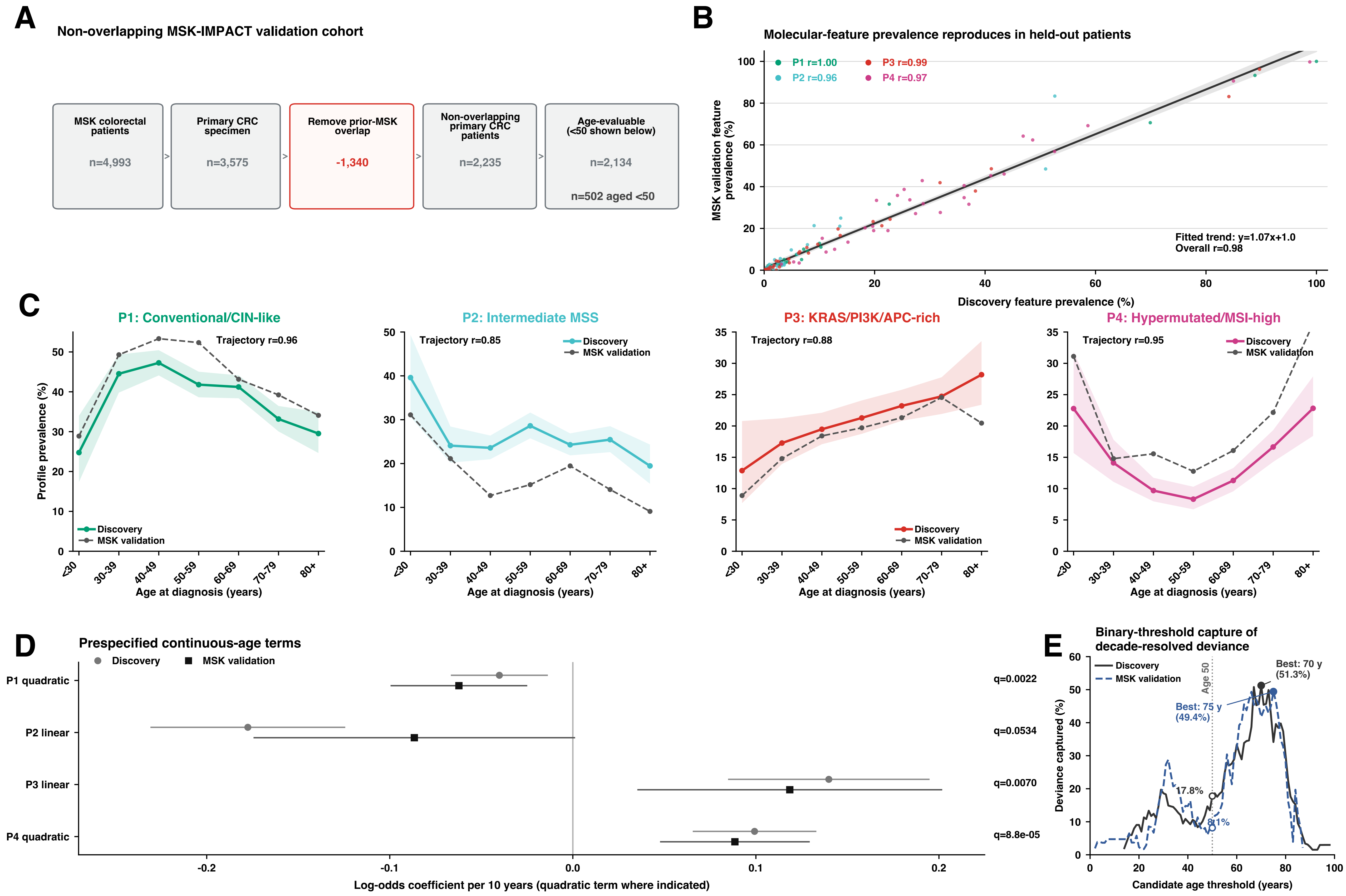
